# Reentrant liquid condensate phase of proteins is stabilized by hydrophobic and non-ionic interactions

**DOI:** 10.1101/2020.05.04.076299

**Authors:** Georg Krainer, Timothy J. Welsh, Jerelle A. Joseph, Jorge R. Espinosa, Sina Wittmann, Ella de Csilléry, Akshay Sridhar, Zenon Toprakcioglu, Giedre Gudiškytė, Magdalena A. Czekalska, William E. Arter, Peter St George-Hyslop, Anthony A. Hyman, Rosana Collepardo-Guevara, Simon Alberti, Tuomas P.J. Knowles

## Abstract

Many cellular proteins demix spontaneously from solution to form liquid condensates. These phase-separated systems have wide-ranging roles in health and disease. Elucidating the molecular driving forces underlying liquid–liquid phase separation (LLPS) is therefore a key objective for understanding biological function and malfunction. Here we show that proteins implicated in cellular LLPS, including FUS, TDP-43, Brd4, Sox2, and Annexin A11, which form condensates at low salt concentrations, can reenter a phase-separated regime at high salt concentrations. By bringing together experiments and simulations, we demonstrate that phase separation in the high-salt regime is driven by hydrophobic and non-ionic interactions, and is mechanistically distinct from the low-salt regime, where condensates are additionally stabilized by electrostatic forces. Our work thus provides a new view on the cooperation of hydrophobicity and non-ionic interactions as non-specific driving forces for the condensation process, with important implications for aberrant function, druggability, and material properties of biomolecular condensates.

## Introduction

Liquid–liquid phase separation (LLPS) has emerged as an important organizing principle in biology, where condensation of proteins and other biomolecules into liquid droplets has been shown to underlie the formation of membraneless subcellular compartments.^1–3^ Beyond compartmentalization, these biomolecular condensates have been implicated in diverse biological processes including chromatin reorganization,^4^ noise buffering,^5^ and sensing^6^, and their misregulation has been associated with the emergence of diverse pathologies, such as neurodegenerative diseases and cancer.^7–9^

LLPS is a thermodynamic process in which interacting multivalent proteins, and in many cases oligonucleotides, minimize their free energy by demixing into a protein-depleted dilute phase and a protein-enriched condensed phase.^10–12^ LLPS becomes thermodynamically favorable at biomolecular concentrations and in solution conditions where the free energy gain from the dynamic formation of weak attractive intermolecular interactions,^13^ and the increase in entropy associated with the release of water molecules from the surfaces of biomolecules to the bulk phase,^14^ become sufficient to overcome the entropy loss due to the reduction in available microstates upon demixing.^10–12^ This intricate balance of entropic and enthalpic forces raises the fundamental question about the nature of the molecular interactions that govern protein LLPS and of the factors that modulate them.

Previous experimental and theoretical studies have shed light on the molecular grammar underlying LLPS.^15–17^ Accordingly, protein condensation has been shown to be driven by the cooperation of both electrostatic and hydrophobic interactions, including charge–charge, cation– π, dipole–dipole, and π–π stacking interactions. The interplay of these interactions underlies the phase separation behavior of proteins at or below physiological ionic strength.

Here we show that a wide range of proteins, including fused in sarcoma (FUS),^18–22^ transactive response DNA-binding protein of 43 kDa (TDP-43),^23,24^ Bromodomain-containing protein 4 (Brd4),^25,26^ Sex determining region Y-box 2 (Sox2),^27^ and Annexin A11 (A11)^28^, which are known to undergo LLPS via homotypic multivalent interactions at low salt concentrations, can also undergo LLPS at high salt concentration (*i.e.*, above 1.5 M NaCl), reentering into a phase-separated regime from a well-mixed state at intermediate salt concentrations. This type of reentrant phase behavior—*i.e.*, where the monotonic variation of a single thermodynamic control parameter drives proteins from a phase-separated state to a macroscopically similar state via two phase transitions^29^—is in contrast to the established RNA-mediated reentrant behavior of protein–RNA coacervates assembled through heterotypic multivalent interactions, which are stable only in the presence of intermediate RNA concentrations, but exist as homogeneous solutions at both high and low RNA concentrations.^30–33^

LLPS at high-salt concentrations has been observed for a few polymer systems and proteins, but only at very high polymer/protein concentrations, typically in the hundreds of micromolar to millimolar range, low temperatures, and/or extremes of pH.^34–43^ Importantly, the reentrant protein LLPS we report here takes place at the low micromolar protein concentrations, temperatures, and pH values typical of physiological LLPS. While occurring at salt concentrations higher than those present physiologically (*i.e.*, at ~2–3 M), the observation of an additional phase transition at high salt underscores the complexity of the dynamic processes that underlie condensate formation and dissolution,^1–3^ and the factors that control them, such as changes in scaffold concentration,^44,45^ fluctuations in the condensate environment,^46,47^ and many others.^32^

Strikingly, our data reveal that the molecular interactions stabilizing condensates in the high-salt reentrant regime are fundamentally distinct from those driving phase separation at low salt. At high salt concentrations, LLPS is mainly driven by hydrophobic and non-ionic interactions. This ability of salt to shift completely the molecular driving forces of protein LLPS is consistent with the wide body of work demonstrating the significance of salt in the modulation of protein stability,^48–51^ protein solubility,^52,53^ protein–protein interactions,^54,55^ and protein–nucleic acid interactions.^56–58^

Hence, our work demonstrates that the preferential interactions that the different amino acids establish in LLPS are heavily context-specific (*i.e.*, they are defined not only by the amino acid chemical makeup but also by the microenvironment and conditions they are exposed to). For example, we show that some amino acid pairs transition from establishing dominant electrostatic attraction or repulsion to engaging instead in strong hydrophobic attraction, as a function of the salt concentration.

As such, this dominant role of hydrophobicity and non-ionic interactions in the high-salt regime expands the molecular grammar governing LLPS, and demonstrates that the driving forces for protein phase separation are not only dictated by the amino acid sequence but also by the condensate environment. Overall, these findings may have wide-ranging implications for the interactions, druggabilities, and material properties of biomolecular condensates, and thus provides a new lens for understanding biomolecular condensate behavior in health and disease.

## Results and Discussion

We first discovered reentrant phase behavior for the protein FUS, when mapping out its phase diagram as a function of KCl concentration (Figure 1). FUS is phase separated in salt concentrations up to ~125 mM, in line with previous observations,^18–22^ and then forms a well-mixed phase between 125 mM and 1.5 M, but surprisingly reenters the phase separated regime above 1.5 M KCl. Hence, FUS exhibits two phase boundaries, at respective upper and lower transition concentrations of salt. Importantly, condensate formation at high KCl concentrations is fully reversible. Adjusting the KCl concentration back to the 500 mM to 1.5 M range yields a well-mixed phase, which is also known to occur for condensates at low salt conditions upon increasing KCl concentration.^21^

**Figure 1.**
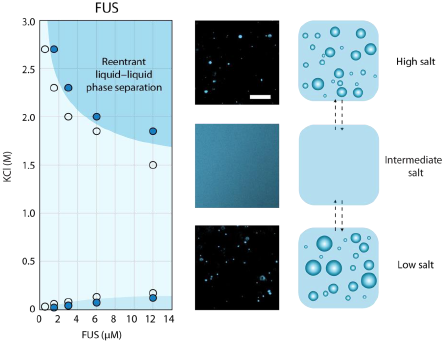
Reentrant phase separation of FUS at high salt. Phase diagram (left), representative images (center), and schematic (right) of FUS phase separation in the presence of increasing concentrations of KCl. In the phase diagram, markers filled with blue indicate concentrations where phase separation was observed in fluorescence images. Open markers indicate concentrations tested where phase separation did not occur. Darker blue regions are guides for the eyes indicating regions where phase separation of FUS occurs, and light blue is the region where no phase separation occurs. The reentrant phase separation regime is indicated. Fluorescent images of FUS (6 μM, EGFP labelled) were taken at 50 mM (low salt), 500 mM (intermediate salt), and 2.7 M KCl (high salt) in 50 mM Tris-HCl (pH 7.2). Scale bar is 20 μm.

In addition to FUS, we find salt-induced reentrant phase behavior in the pathological G156E mutant of FUS,^18,19^ TDP-43,^23,24^ Brd4,^25,26^ Sox2,^27^ and A11^28^ (Figure 2). Like FUS, all these proteins phase-separate via homotypic multivalent interactions at low salt, are involved in essential cellular and developmental processes,^59,60^ and implicated in neurodegenerative disorders.^7,28^ Notably, in all cases, high-salt condensates have similar size distributions and shapes as their low-salt counterparts. An analysis of droplet shape revealed that both low- and high-salt condensates exhibit a high degree of circularity (>95%), and have similar areal distributions (see Supplementary Information, Figure S1), substantiating their liquid-like character and structural similarities in both salt regimes.

**Figure 2.**
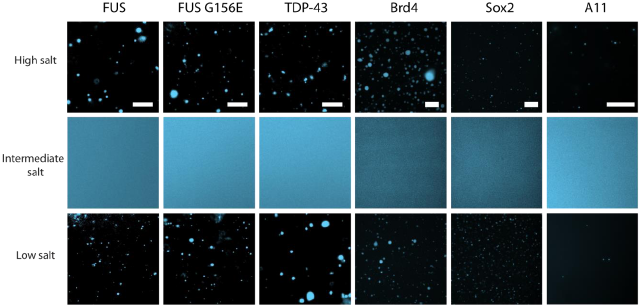
Salt-mediated reentrant phase separation of FUS, FUS G156E, TDP-43, Brd4, Sox2, and A11. Representative images of FUS, FUS G156E, TDP-43 at 50 mM (low salt), 500 mM (intermediate salt), and 2.7 M KCl (high salt) in 50 mM Tris-HCl (pH 7.2); Brd4 and Sox2 at 50 mM (low salt), 500 mM (intermediate salt), and 2.15 M KCl (high salt); Brd4 buffer: 5 mM Tris (pH 7.5), 0.2 mM EDTA, 0.5% glycerol; Sox2 buffer: 5 mM Bis-Tris-Propane (pH 7.5), 0.5% glycerol; and A11 at 22.5 mM (low salt), 225 mM (intermediate salt), and 500 mM NaCl (high salt) in 20 mM HEPES (pH 7.0). For fluorescence imaging, both FUS variants and TDP-43 were tagged with EGFP and studied at a protein concentration of 6 μM; Brd4 and Sox2 were tagged with monoGFP and studied at protein concentrations of 6 μM and 12.4 μM, respectively; A11 was labelled with AlexaFluor647 and studied at a protein concentration of 15 μM. Scale bars are 20 μm in all images.

The observed reentrant phase behavior provides an opportunity to shed light on the molecular processes that lead to condensate formation in the high-salt regime, and to probe whether these phenomena are the same at high and low salt concentrations. Condensation at and below physiological salt concentrations, for example of FUS,^18,19^ is mainly driven by an interplay of both electrostatics^21,61^ and hydrophobic interactions,^62,63^ and can be modulated by RNA^20^ and ATP^64,65^ both of which favor and disrupt phase separation depending on their concentration. This broad mix of forces that drive and modulate phase separation at physiological salt concentrations is supported by the widespread use of charge-modifying post-translational modifications in cells to control the strength of protein–protein^66^ and protein–DNA interactions,^67^ and the ability of uncharged proteins to fold and self-assemble into large complexes.^68^

From these observations it follows that the molecular processes that lead to demixing and condensate formation in the high-salt regime are likely connected to electrostatic protein–protein interactions becoming negligible and the hydrophobic effect being heightened. Indeed, the significant drop in protein solubility upon the addition of salt, which can result in precipitation for many proteins (*i.e.*, the ‘salting out’ effect), has been attributed to hydrophobic effects.^48,69^ Thus, we reasoned that enhanced hydrophobic interactions and weakened ionic interactions might be the key drivers of protein reentrant phase separation at high-salt concentration.

To test this hypothesis, we probed the ability of pre-formed FUS condensates to dissociate when exposed to a range of additional components acting as phase separation disruptors, as shown in Figure 3a. We selected a representative set of compounds with the ability to modulate both electrostatic interactions, such as poly-uridine (PolyU) RNA and ATP, both highly negatively charged molecules previously described to disrupt phase separation,^20,64,65^ as well as 1,6-hexanediol, an aliphatic alcohol known to disrupt weak protein–protein hydrophobic interactions and selectively dissolve liquid condensates but not solid ones.^70^ At low salt concentrations, PolyU RNA, ATP, and 1,6-hexanediol were all able to dissolve FUS condensates, confirming that both hydrophobic and electrostatic interactions contribute to the stability of FUS condensates in the low-salt regime. At high-salt conditions, 1,6-hexanediol was the only disruptor that could dissolve FUS condensates, while addition of PolyU RNA and ATP did not show any effects. These observations, summarized in Figure 3b, suggest that reentrant high-salt phase separation of proteins is indeed primarily a hydrophobically driven process where electrostatics are screened out.

**Figure 3.**
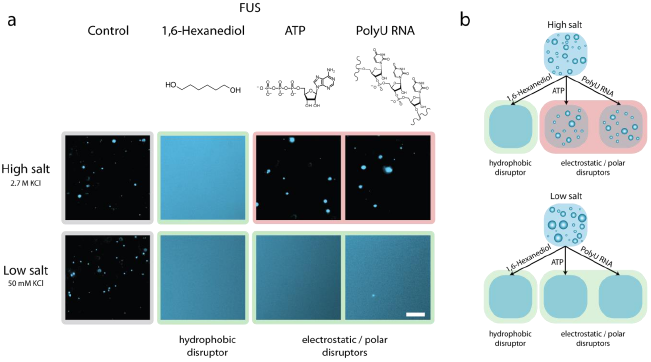
Dissolution assay of FUS condensates in the high- and low-salt regime using hydrophobic and electrostatic/polar disruptors. **(a)** Representative images of FUS condensates upon addition of 1,6-hexanediol, ATP, and PolyU RNA are shown. Total protein concentration was 4.5 μM and final additive concentrations were 10% 1,6-hexanediol, 1.25 mg/mL PolyU RNA, 12.5 mM ATP in 50 mM Tris-HCl (pH 7.2) at 50 mM (low salt) and 2.7 M KCl (high salt). Conditions at which the disrupters dissolved the condensates are highlighted in green and those where condensates remained intact are highlighted in red. Scale bar is 20 μm. **(b)** Schematic representation of the ability for electrostatic/polar disruptor molecules ATP and PolyU RNA to dissolve condensates in the low-salt regime but not in the high-salt regime, and for the hydrophobic disruptor 1,6-hexanediol to dissolve condensates in both regimes.

To understand how modulation of salt concentration impacts phase behavior more generally, we probed the response of the highly positively-charged PR_25_ peptide, which is formed by 25 repeats of the proline–arginine dipeptide (Figure 4a). Unlike FUS, PR_25_ does not exhibit LLPS at low salt concentrations because, in this regime, its homotypic interactions are dominated by the Arg–Arg repulsion; instead PR_25_ requires the addition of co-factor polyanions such as RNA to form complex coacervates at physiological salt.^71,72^ However, and strikingly, at KCl concentrations of 2.7 M, we find that PR_25_ undergoes LLPS on its own (Figure 4a). Similar to FUS in the high-salt regime, 1,6-hexanediol could dissolve these high-salt PR_25_ condensates, while the addition of PolyU RNA and ATP did not elicit any observable effects (Figure 4b). These results suggest that phase separation of PR_25_ at high salt is also driven by hydrophobic interactions. Notably, this type of phase transition is not an example of reentrant phase behavior, yet it provides an additional demonstration of the occurrence of homotypically-driven LLPS at high salt, enabled by the screening of electrostatic interactions, in this case repulsion among Arg residues, and strengthening of non-charged interactions.

**Figure 4.**
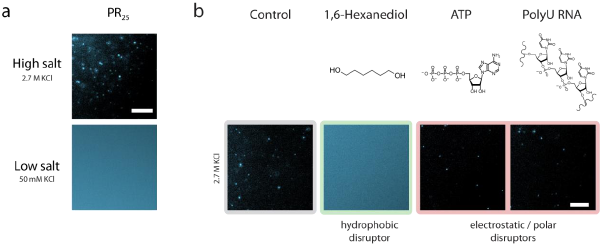
Phase separation and disruptor-mediated dissolution behavior of the PR25 peptide at high and low salt concentrations. **(a)** Representative images of PR_25_ at 50 mM (low salt) and 2.7 M KCl (high salt) in 50 mM Tris-HCl (pH 7.2). Unlabeled peptide was mixed with a small amount of the same peptide labeled with AlexaFluor546; total peptide concentration was 72 μM. **(b)** Dissolution assay of PR_25_ condensates in the high-salt regime using hydrophobic (1,6-hexanediol) and electrostatic/polar disruptors (ATP and PolyU RNA). Final peptide concentration was 54 μM PR_25_ and final additive concentrations were 10% 1,6-hexanediol, 1.25 mg/mL PolyU RNA, 12.5 mM ATP in 2.7 M KCl, 50 mM Tris-HCl (pH 7.2). Conditions at which the disruptors dissolved the condensates are highlighted in green and those where condensates remained intact are highlighted in red. Only 1,6-hexanediol dissolves PR_25_ condensates at high salt. Scale bars in all images are 20 μm.

To explore further the role of hydrophobicity in phase separation in the high-salt regime, we systematically varied the chemical nature of the salts used along the Hofmeister series,^73,74^ as was previously done for the low-complexity domain of FUS at lower salt concentrations.^62^ The Hofmeister series is ordered based on the ability of ions to reduce the solubility of hydrophobic molecules in water (*i.e.*, the ‘salting-out’ effect). Although the exact molecular-level mechanism explaining the salting-out effect remains controversial,^75^ it has been suggested that salts higher up in the Hofmeister series increase the solubility of proteins in solution by effectively weakening the strength of hydrophobic interactions. Therefore, ascending the Hofmeister series, higher salt concentrations should be required to induce phase separation. To test this hypothesis, we mapped out the phase behavior of FUS and PR_25_ in the high-salt regime using chloride salts of various Hofmeister series cations (Figure 5a,b). Indeed, the phase boundary shifts in the order as given by the most agreed upon ranking of the series^76^ (*i.e.*, K^+^ < Na^+^ < Rb^+^ < Cs^+^ < Li^+^ < Ca^2+^); hence, condensate formation is disfavored with salts higher up in the series. Similarly, for both FUS and PR_25_, less 1,6-hexanediol is required to dissolve condensates made in solutions containing salts higher up in the Hofmeister series (Figure 5c,d). These observations support the hypothesis that the formation of FUS and PR_25_ condensates at high salt concentration is driven by hydrophobic interactions, and is thus of a different nature to the transition observed at low salt concentrations, which has a significant electrostatic contribution. Our results also demonstrate that, analogous to the salting-out effect, salts across the Hofmeister series have a different impact on modulating the solubility limit of phase-separating proteins, and in shifting the boundary of immiscibility that determines phase separation.

**Figure 5.**
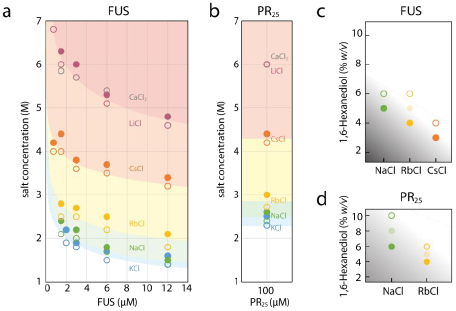
Hofmeister effect in the high-salt phase separation behavior of FUS and PR_25_. **(a)** Phase diagram for FUS as a function of salt concentration of various salts of the Hofmeister series. Open circles indicate cases where phase separation did not occur, closed circles indicate where phase separation did occur. Each curve depicts the apparent phase boundary for the particular salt named next to it and is only meant as a guide for the eyes. Even at the saturation concentration of CaCl_2_ (grey), the hydrophobic effect is weakened to the extent such that phase separation cannot occur, indicated by the presence of open circles and absence of closed ones in the phase diagram. **(b)** Phase behavior of PR_25_ as a function of ionic strength of various salts. The trend is consistent with panel a. **(c)** Comparison of the amount of 1,6-hexanediol required to dissolve FUS condensates in solutions of various salts along the Hofmeister series. In each solution, the final salt concentration was 4 M and the final FUS concentration was 2 μM. Partially shaded circles represent conditions where the number of condensates was visibly reduced, but the condensates were not fully dissolved. **(d)** Comparison of the amount of 1,6-hexanediol required to dissolve PR_25_ as a function of salts along the Hofmeister series. The final salt concentration at each point was 4 M and the PR_25_ concentration was 100 μM.

Next, we investigated the molecular driving forces behind reentrant protein phase behavior, by developing a multiscale modeling approach that combines molecular dynamics (MD) simulations at two complementary levels of resolution: atomistic simulations of amino acid pairs and residue-resolution coarse-grained simulations of protein condensates (see Supplementary Information). This combination of simulations at two levels of resolution allows us to investigate how the relative contributions of electrostatic and hydrophobic interactions among atomistic amino acid pairs change as a function of salt, and subsequently to determine if such changes are consistent with protein reentrant phase behavior.

To this end, we first performed a set of all-atom umbrella sampling MD simulations of amino acid pairs combining residues with different chemical makeups (*i.e.*, charge, polar atoms, *sp^2^*-hybridized atoms, and aromatic groups) using the state-of-the-art AMBERff03ws force field^77^ and explicit solvent and ions. From these simulations, we extract the potential of mean force (PMF) as a function of the center-of-mass (COM) distance between amino acids at different salt concentrations. By monitoring the salt-dependent changes to the various PMF minima, we estimated how the free energy that stabilizes the bound configuration of an amino acid pair is modulated by salt and the chemical composition of the pair. Since we are interested in probing interaction potentials up to very high salt concentrations (*i.e.*, from 0 to 3 M NaCl), it is important that the solvent and ion model parameters employed are well-fitted to reproduce adequately ion solubilities in water at 298 K (*i.e.*, the absence of unphysical salt crystallization). We used the JC-SPC/E-ion/TIP4P/2005 force field, which has been optimized for that purpose, and verified that the correct ion solubilities are observed.^78^

Because of their dependency on polarization, investigating atomistically the effects of salt on cation–π interactions is not trivial. Cation–π interactions at low salt involve the electrostatic attraction between the polarizable quadrupole of the π-electron cloud of an aromatic ring and a polarizing positively charged amino acid.^79^ Fixed charge atomistic force fields ignore polarization effects and are, therefore, unable to properly capture cation–π interactions.^80,81^ While approximations to incorporate the many-body effects of polarization in atomistic force fields exist, these require many iterations at each molecular configuration, which comes at a huge computational cost, and are not free from their own inaccuracies.^82^ Furthermore, the polarizing power of the cation is screened out as the salt concentration increases, which implies that the cation–π pair is significantly polarized only at low salt. Therefore, to study cation–π interactions in the context of salt-dependent protein LLPS, we have developed a specialized model by refitting the sidechain charges of Tyr and Phe amino acids when bound to Arg or Lys (at the MP2/6-31G(d) level of theory) to describe the post-polarized state of the pairs in the low-salt regime, and assumed no polarization in the moderate and high-salt regimes, where significant cation screening is expected (see Supplementary Information).

The PMF curves (Figure 6) show that the attractive interaction energies at short inter-molecular distances among oppositely charged amino acids decrease monotonically with increasing NaCl concentration (Figure 6a,b). Conversely, those involving only uncharged residues, including polar ones, increase significantly with salt (*i.e*., Ala–Ala, Pro–Pro, Ser–Ser, and Tyr–Tyr) (Figure 6c–f). Notably, interactions among a basic and an aromatic residue (*e.g.*, Arg–Tyr or Lys–Phe, typically termed cation–π) exhibited a more complex hybrid (electrostatic– hydrophobic) behavior (Figure 6g; discussed below), which yields a high interaction strength at both low and high salt. Likewise, Arg–Arg (Figure 6h) present a switch-like behavior and become attractive at high salt. A summary is given in Figure 6i. In all cases, the strongest attractive interactions we observe, both at low and high salt, are those where the two amino acids in the pair have π-electrons (*e.g.*, Arg–Glu, Tyr–Tyr and Arg–Tyr/Phe) (see Figure S2). Together, these observations support our hypothesis that, at low salt, protein condensation is stabilized by a combination of electrostatic and non-ionic forces, while at high salt the hydrophobic contributions are strongest, and further suggest a dominant role of π–π interactions in both regimes.

**Figure 6.**
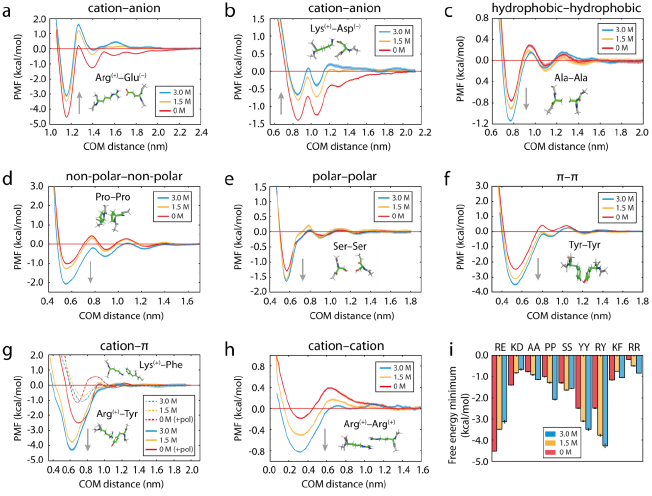
Effect of salt (0 M, 1.5 M, 3 M NaCl) on the potential of mean force (PMF) between selected amino acid pairs in explicit solvent and NaCl ions as a function of the center-of-mass (COM) distance. **(a)** cation–anion (with π–π contributions), **(b)** cation–anion (without π–π contributions), **(c)** hydrophobic–hydrophobic, **(d)** non-polar– non-polar, **(e)** polar–polar, **(f)** π–π, **(g)** hybrid cation–π/π–π (Arg–Tyr, solid lines) and cation–π (Lys–Phe, dashed lines) (+pol denotes refitted Tyr/Phe parameters were employed; as described in the text and the Supplementary Information), **(h)** cation–cation (with π–π contribution). The second well in (b) emerges from the interaction of Asp with an additional H atom in the Lys sidechain, which is displaced by about 1.7 Å from the two H atoms that contribute to the first well. To evaluate (g), a model for the polarized cation–π systems was developed (see Methods). The grey arrows in each panel highlight the general shift direction of the PMF minimum as salt concentration is raised. Upward arrows show weakening of cation–anion interactions upon increasing salt. Downward arrows show strengthening of non-ionic interactions and of hybrid cation–π/π–π and cation–cation interactions when both amino acids in the pair have π–orbitals. Error bars are shown as bands and represent the standard deviations obtained by bootstrapping the results from three independent simulations. **(i)** Variation in the free energy minimum for profiles in a–h (with associated error bars) with salt. One-letter amino-acid codes are used to identify each pair interaction.

Although interactions among cationic amino acids and aromatic residues are commonly described as mostly electrostatic—with polarization forces playing a significant role^83^—we find that they are less sensitive to screening than interactions between oppositely charged amino acids (Figure S2f). This is consistent with experimental measurements of peptide helicity showing that cation–aromatic interactions are not fully disrupted by neutral salts up to concentrations of 2.5 M.^84,85^ High-level quantum chemical calculations attribute the lower sensitivity of cationic and aromatic pairs to screening to their higher orbital coordinate contributions.^86^ Indeed, our results support the hypothesis interactions between basic amino acids and aromatic rings switch from cation–π electrostatic in nature at low salt to hydrophobic at high salt and are, therefore, stabilized.

Despite it being universal, the stabilization of hydrophobic cation–aromatic interactions at high salt has diverse molecular origins that depend on the exact chemical makeup of each pair. Lys–Phe is an example of a pair that establishes a more pure cation–π bond at low salt in which no π-electrons are contributed from the Lys cation sidechain. This interaction first decreases significantly from low to moderate salt, alluding to its significant electrostatic character (Figure 6g; dashed lines). However, contrary to a solely electrostatic interaction, the Lys–Phe interaction has a shorter range and, in the high salt regime, it regains its low-salt attractive power. This occurs because, at high salt, the π-electrons in the aromatic ring of Phe interact preferentially and more strongly with the methyl groups of Lys.^87^

Most strikingly, and in contrast to the Lys–Phe cation–π pair, our PMFs reveal that the Arg–Tyr/Phe bond is better described as a strongly attractive hybrid cation–π/π–π interaction that increases monotonically with salt (Figure 6g; solid lines and Figure S2). Both the significant strengthening with salt and the high magnitude of the Arg–Tyr/Phe interaction throughout are explained by a strong contribution of π–π bonding between the π–orbitals in the *sp*^2^-hybridized guanidinium group of Arg and the π–electrons of the aromatic rings.^87–90^ This important π–π contribution was further demonstrated by performing additional PMF calculations in which the sidechain π–orbitals of Arg and Tyr were maintained in a perpendicular (t-shaped) arrangement that does not favor π–π bonding (Figure S3c). In contrast to the parallel Arg–Tyr pair interaction (which includes π–π stacking and, therefore, increases monotonically with increasing salt), the variation of the t-shaped Arg–Tyr interaction with salt mirrors that of Lys–Phe and is much weaker than its parallel counterpart (Figure S3e). Moreover, for Lys–Phe the interaction is negligibly impacted by the pair conformation (Figure S3b and d), which further underscores the dominant role of π–π bonding in Arg–Tyr/Phe cation–aromatic interactions, but not in Lys–Phe.

The crucial role of π–π bonds as a driving force for protein LLPS across all salt regimes, but particularly at high salt, is evident not only from the deep wells in the PMFs of *sp*^2^-hybridized cation–aromatic and aromatic–aromatic pairs (Figure 6i), but also from the behavior of charge– charge pairs with π electrons. A striking example of this is the transition of the interaction among Arg–Arg from mainly an electrostatic repulsive interaction at low salt to a weak attractive π–π interaction^89,91^ in the high-salt regime (Figure 6h). This transition occurs because once the repulsion among positive guanidinium groups is screened, the *sp*^2^-hybridized planar guanidiniums can interact via their π–orbitals. Glu, Asp, and nucleotides also have charged groups and π-orbitals, and hence, their homotypic interaction could similarly exhibit this striking transition from repulsion to attraction as salt increases. Moreover, π–π bonding between Arg and nucleotides is consistent with the much higher salt-range stability observed experimentally for droplets co-assembled with poly-Arg and polynucleotides over those formed with poly-Lys and polynucleotides.^90^ These unexpected results further illuminate the differences in the molecular interactions stabilizing protein condensates at high salt versus low salt and put forward Arg–Arg as an additional force that stabilizes the homotypic LLPS of PR_25_ in the high-salt regime. We therefore propose that at ionic strengths where charges are screened, π-driven hydrophobic interactions may sustain LLPS.

In a next step, using a coarse-grained modelling approach, we investigated whether salt-modulation of intermolecular interactions, observed atomistically, is indeed responsible for protein LLPS at high salt. For this purpose, we conducted direct coexistence simulations of tens of interacting FUS and PR_25_ polypeptide chains using a reparameterization of the amino-acid resolution coarse-grained model of the Mittal group,^16,92,93^ which considers sequence-dependent electrostatic and hydrophobic interactions; we developed this reparameterization to recapitulate the higher experimental LLPS propensity at low salt of FUS over its prion like domain^94^ (see Supplementary Information). To investigate salt-dependent LLPS of FUS and PR_25_ (Figure 7a), we modulated the relative contribution of electrostatic and hydrophobic interactions among amino acid pairs in the coarse-grained model, according to the salt concentration, based on our atomistic results (Figure 6i). Consistent with our experiments at low salt, we observe LLPS for FUS (due to strong attractive electrostatic cation–anion and cation–π interactions), but not for PR_25_, which is highly enriched in positively charged amino acids that repel each other strongly in this regime (Figure 7b, all interactions). To confirm the dependence of LLPS on electrostatic forces at low salt, we scaled down the charged–charged interactions (as suggested by our PMFs) and, as expected, observed melting of the FUS condensates (Figure 7c; reduced electrostatics); this finding corroborates the key role of electrostatic interactions in stabilizing protein condensates at low salt. Finally, to recapitulate reentrant phase behavior at high salt, we moderately increased the strength of the hydrophobic interactions (by only 10% for FUS, including the hydrophobic attraction from the cation–π pairs at high salt, and 30% for PR_25_, as suggested by our PMF calculations), while keeping the electrostatic interactions scaled down. Indeed, we observed that this subtle enhancement of hydrophobic attraction is sufficient to yield a reentrant phase transition for FUS and induced phase separation for PR_25_ (Figure 7d; reduced electrostatics + increased hydrophobicity). Overall, these results show that protein LLPS in the high-salt regime is driven by hydrophobic interactions with the strongest contribution coming from π–π bonds; thereby, providing a molecular explanation for our experimental observations. Our simulations further demonstrate that despite the differences in the molecular driving forces stabilizing FUS condensates at low and high salt, their molecular organization (Figure S4) and densities (Table S3) are similar in both regimes. Hence, by approaching either extremes of salt, the system does in fact reenter a previously encountered phase-separated state.

**Figure 7.**
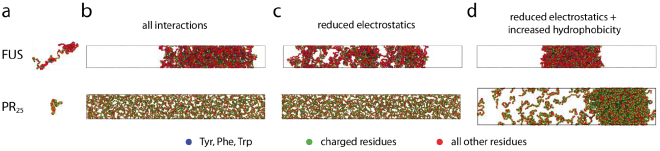
Dependence of LLPS on electrostatic versus hydrophobic forces for FUS and PR25 from direct coexistence simulations using a sequence-dependent protein coarse-grained model. **(a)** Illustration of the coarse-grained models for the different proteins with one bead representing each amino acid. Amino acids are colored according to their chemical identity (aromatics in blue, charged residues in green, all other residues in red; color code shown at the bottom). Snapshots for simulations with **(b)** ‘all interactions’, **(c)** ‘reduced electrostatics’, and **(d)** ‘reduced electrostatics + increased hydrophobicity’ for FUS (24 proteins) and PR_25_ (400 peptides).

## Conclusion

This work demonstrates the existence of reentrant protein phase separation as a function of salt concentration as the control parameter. Our results show that protein molecules are driven by homotypic multivalent interactions to demix from a homogeneous phase in the limits of both low and high electrostatic screening. Previously, a different type of reentrant phase transition from the well-mixed to the phase-separated and back to the well-mixed state has been described for systems that phase separate via heterotypic protein–RNA interactions at intermediate RNA concentrations exclusively.^30–33^ Additional theoretical and experimental work has predicted the ability for reentrant phase separation of proteins, peptides, and polymers to occur as a function of pH,^95^ temperature,^93,96,97^ and pressure.^98,99^ Our work presents a novel type of reentrant phase transition in which proteins can phase separate on their own in two distinct regimes in response to changes of ionic strength. Analysis of the nature of the molecular interactions implicated in biomolecular phase transitions shows that phase separation in the absence of charge screening at low salt concentration is driven by the cooperation of electrostatic and hydrophobic interactions, while the same process at high salt concentration is favored mainly by hydrophobic and non-ionic interactions, such as interactions between Ala–Ala, Pro–Pro, Tyr–Tyr, Ser–Ser, Arg–Tyr, Arg– Arg. The latter two, as we have shown in this study, become predominantly hydrophobic in the high salt regime and interact under those conditions predominantly through π–π bonds (Figure 8). Importantly, we show that π–π interactions, involving both aromatic and non-aromatic residues, are dominant driving forces for LLPS in both salt regimes, but most strongly under high salt conditions. Our work thus provides a new view of the role of hydrophobicity and non-ionic interactions as non-specific driving forces for the condensation process and expands the molecular grammar of interactions governing LLPS of proteins.

**Figure 8.**
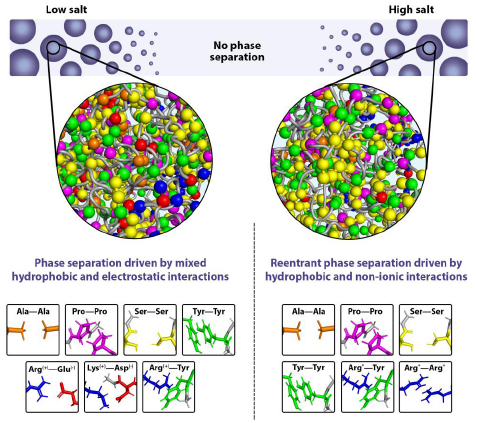
Schematic illustration of the different molecular forces that stabilize condensates in the low-salt versus the high-salt reentrant regime. While phase separation in the low-salt regime is driven by both electrostatic and hydrophobic interactions, the condensation process in the reentrant high-salt regime is governed by hydrophobic and non-ionic interactions. Note: The asterisks (*) for Arg*–Try and Arg*–Arg* indicate that at high salt, charges are screened, and interactions become predominantly hydrophobic (*i.e.*, π–π interactions).

The observed salt-mediated modulation of LLPS underscores the finding that the molecular driving forces for phase separation are not only dictated by the protein sequence but are also crucially sensitive to solution conditions. Solution conditions are essential to LLPS because they regulate the competition between the free-energy reduction stemming from favorable protein– protein interactions in conjunction with the release of water molecules to the bulk phase, and the entropic cost of demixing. Hence, solution conditions modulate the preferential interactions among amino acids.^1,14^ Strikingly, our work shows that the Arg–Arg and Lys–Phe pairs exhibit a salt-dependent switch-like behavior where they transition from establishing a dominant electrostatic interaction (cation–cation repulsion and cation–π attraction, respectively) at low salt to a hydrophobic attraction at high salt stabilized by a strong π–π component (for Arg–Arg) or methyl– π interaction (for Lys–Phe). The tuneability with salt therefore highlights the complex interplay of specific intermolecular interactions that commit proteins to particular phases with the condensate environment, and demonstrates the ability for reentrant phase transitions to occur in biomolecular systems under conditions of varying electrostatic screening.

The finding of the existence of salt-dependent reentrant phase separation provides a new view on the cooperation of hydrophobicity and non-ionic interactions as non-specific driving forces for the condensation process, and may have wide-ranging implications for the druggabilities and material properties of biomolecular condensates. For example, aberrant phase separation of FUS, TDP-43, and Annexin A11 has been associated with neurodegenerative diseases.^7^ The discovery of salt-mediated reentrant phase separation for these systems suggests a crucial feature of protein LLPS behavior that may be important when developing therapies for phase-separation implicated diseases.^100^ Hence, drugs that are designed to prevent or reverse phase separation, yet induce high electrostatic screening, could in turn trigger reentrant transitions. Thus, therapies must be aimed at transitioning pathological condensates into an intermediate and not a high electrostatic screening regime.

Moreover, pathological liquid-to-solid phase transitions in FUS, for example, are thought to be linked to the emergence of strongly-bound ladders of paired kinked β-sheets, which assemble by a combination of hydrogen bonding and predominant π–π hydrophobic interactions among aromatic residues within “low-complexity aromatic-rich kinked segments” (LARKS) that assemble into protofilaments.^101^ FUS-derived peptides that contain LARKS and are devoid of charged residues interact weakly under physiological conditions (*i.e.*, salt and temperature), as shown by their phase-separation being enhanced at higher salt concentrations.^33^ This observation suggests that, under physiological conditions, LARKS-driven pathological transitions are disfavored. However, in the reentrant high-salt regime, or conditions in cells that give rise to similar electrostatic screening (*e.g.*, the presence of multivalent ions or other charged species and post-translational chemical modifications), our work suggests that pathological LARKS–LARKS interactions would be significantly enhanced due to strengthening of π–π interactions, thereby enabling pathological transitions.

Taken together, our study identifies a novel salt-mediated reentrant protein LLPS behavior that is enabled by hydrophobic and non-ionic interactions. The discovery of high-salt protein phase separation provides a compelling view of the plasticity of the molecular driving forces for protein phase separation, and emphasizes that these forces are not only defined by the amino acid sequence, but also critically influenced by the condensate environment. Depending on the microenvironment, the diverse chemical makeups of amino acids allows them to engage in a multiplicity of molecular interactions (*e.g.*, hydrophobic, electrostatic, mixed), going beyond— hence expanding—the typical one-way classification they are traditionally given (*e.g.*, polar, hydrophobic, charged). These findings highlight the importance of considering solution conditions to which condensates are exposed when aiming at predicting, rationalizing, or modulating protein phase behavior, and when designing therapies to ameliorate phase separation-related pathologies.

## Methods

Details on experimental materials and methods, and molecular modelling and simulation methods are available in the Supplementary Information.

## Data availability

The authors confirm that all data generated and analyzed during this study are included in this article and its Supplementary Information. Data are also available from the corresponding authors upon request.

## Code availability

Computer code and data analysis codes used in this article and its Supplementary Information are available from the corresponding authors upon request.

## Acknowledgments

We thank Rohit V. Pappu for helpful comments and stimulating discussions. We thank Jeetain Mittal and Gregory L. Dignon for invaluable help with the implementation of their sequence-dependent protein coarse-grained model in LAMMPS. The research leading to these results has received funding from the European Research Council (ERC) under the European Union’s Seventh Framework Programme (FP7/2007-2013) through the ERC grant PhysProt (agreement no. 337969) (T.P.J.K.), under the European Union’s Horizon 2020 Framework Programme through the Future and Emerging Technologies (FET) grant NanoPhlow (agreement no. 766972) (T.P.J.K., G.K.), the Marie Skłodowska-Curie grant MicroSPARK (agreement no. 841466) (G.K.), the Marie Skłodowska-Curie grant StressGranule (agreement no. 791147) (S.W.), and the ERC grant InsideChromatin (agreement no. 803326) (R.C.-G.). We further thank the Newman Foundation (T.P.J.K.), the Biotechnology and Biological Sciences Research Council (T.P.J.K.), the Herchel Smith Funds (G.K.), the Wolfson College Junior Research Fellowship (G.K.), the Winston Churchill Foundation of the United States (T.J.W.), the Harding Distinguished Postgraduate Scholar Programme (T.J.W.), the Winton Advanced Research Fellowship (R.C.-G.), the King’s College Research Fellowship (J.A.J.), the Oppenheimer Research Fellowship (J.R.E.), and Emmanuel College Roger Ekins Fellowship (J.R.E). M.A.C. was supported by the Polish Ministry of Science and Higher Education within the Mobilność Plus V fellowship (decision number 1623/MOB/V/2017/0). We also acknowledge funding from the Canadian Institutes of Health Research (Foundation Grant and Canadian Consortium on Neurodegeneration in Aging Grant) (P.StG.-H.), the Wellcome Trust Collaborative Award 203249/Z/16/Z (P.StG.-H., T.P.J.K.), the ALS Canada Project Grant and the ALS Society of Canada/Brain Canada (grant no. 499553 (P.StG.-H.), the Alzheimer’s Research UK (ARUK) and the Alzheimer’s Society UK (P.StG.-H.), and the US Alzheimer Society Zenith Grant ZEN-18-529769 (P.StG.-H.). The simulations were performed using resources provided by the Cambridge Tier-2 system operated by the University of Cambridge Research Computing Service (http://www.hpc.cam.ac.uk) funded by EPSRC Tier-2 capital grant EP/P020259/1.

## Author contributions

Conceptualization, G.K., T.J.W., T.P.J.K.; Formal Analysis, J.A.J., J.R.E., A.S.; Funding acquisition, G.K., T.J.W., J.A.J., J.R.E., S.W., M.A.C., P.StG.-H., R.C.-G., S.A., T.P.J.K.; Investigation, G.K., T.J.W., J.A.J., J.R.E., R.C.G., S.W., E.C., A.S., M.A.C.; Methodology, G.K., T.J.W., J.A.J., J.R.E., R.C.-G., T.P.J.K.; Project administration, G.K., R.C.-G., T.P.J.K.; Resources, T.J.W., P.StG.-H., A.A.H., S.A.; Software, J.A.J., J.R.E; Supervision, G.K., R.C.-G., S.A., T.P.J.K.; Visualization, G.K., T.J.W., J.A.J., J.R.E., E.C., M.A.C.; Writing – original draft, G.K., T.J.W., R.C.-G., J.A.J.; Writing – review & editing, G.K., T.J.W., R.C.-G., J.A.J., J.R.E., S.W., E.C., A.S., Z.T., G.G., M.A.C., W.E.A., P.StG.-H., A.A.H., S.A., T.P.J.K.

## Additional information

**Supplementary Information** is available for this paper: Methods (Materials, Protein production, Sample preparation and generation of phase diagrams, Fluorescence imaging, Simulation Methods), Supplementary Information for atomistic PMF calculations (Tables S1 and S2), Supplementary Information for condensate densities (Table S3), Supplementary Figures (Figures S1, S2, S3, S4), Supplementary References.

## References

1. Hyman, A. A., Weber, C. A. & Jülicher, F. Liquid-Liquid Phase Separation in Biology. Annu. Rev. Cell Dev. Biol. 30, 39–58 (2014).

2. Banani, S. F., Lee, H. O., Hyman, A. A. & Rosen, M. K. Biomolecular condensates: organizers of cellular biochemistry. Nat. Rev. Mol. Cell Biol. 18, 285–298 (2017).

3. Shin, Y. & Brangwynne, C. P. Liquid phase condensation in cell physiology and disease. Science 357, eaaf4382 (2017).

4. Welsh, T. J., Shen, Y., Levin, A. & Knowles, T. P. J. Mechanobiology of Protein Droplets: Force Arises from Disorder. Cell 175, 1457–1459 (2018).

5. Klosin, A. et al. Phase separation provides a mechanism to reduce noise in cells. Science 367, 464–468 (2020).

6. Yoo, H., Triandafillou, C. & Drummond, D. A. Cellular sensing by phase separation: Using the process, not just the products. J. Biol. Chem. 294, 7151–7159 (2019).

7. Alberti, S. & Dormann, D. Liquid–Liquid Phase Separation in Disease. Annu. Rev. Genet. 53, 171–194 (2019).

8. Molliex, A. et al. Phase Separation by Low Complexity Domains Promotes Stress Granule Assembly and Drives Pathological Fibrillization. Cell 163, 123–133 (2015).

9. Bouchard, J. J. et al. Cancer Mutations of the Tumor Suppressor SPOP Disrupt the Formation of Active, Phase-Separated Compartments. Mol. Cell 72, 19–36.e8 (2018).

10. Berry, J., Brangwynne, C. P. & Haataja, M. Physical principles of intracellular organization via active and passive phase transitions. Reports Prog. Phys. 81, 046601 (2018).

11. Bentley, E. P., Frey, B. B. & Deniz, A. A. Physical Chemistry of Cellular Liquid‐Phase Separation. Chem. – A Eur. J. 25, 5600–5610 (2019).

12. Dignon, G. L., Best, R. B. & Mittal, J. Biomolecular Phase Separation: From Molecular Driving Forces to Macroscopic Properties. Annu. Rev. Phys. Chem. 71, 53–75 (2020).

13. Brangwynne, C. P., Tompa, P. & Pappu, R. V. Polymer physics of intracellular phase transitions. Nat. Phys. 11, 899–904 (2015).

14. Ribeiro, S. S., Samanta, N., Ebbinghaus, S. & Marcos, J. C. The synergic effect of water and biomolecules in intracellular phase separation. Nat. Rev. Chem. 3, 552–561 (2019).

15. Wang, J. et al. A Molecular Grammar Governing the Driving Forces for Phase Separation of Prion-like RNA Binding Proteins. Cell 174, 688–699 (2018).

16. Dignon, G. L., Zheng, W., Kim, Y. C., Best, R. B. & Mittal, J. Sequence determinants of protein phase behavior from a coarse-grained model. PLOS Comput. Biol. 14, e1005941 (2018).

17. Alberti, S. Phase separation in biology. Curr. Biol. 27, R1097–R1102 (2017).

18. Patel, A. et al. A Liquid-to-Solid Phase Transition of the ALS Protein FUS Accelerated by Disease Mutation. Cell 162, 1066–1077 (2015).

19. Murakami, T. et al. ALS/FTD Mutation-Induced Phase Transition of FUS Liquid Droplets and Reversible Hydrogels into Irreversible Hydrogels Impairs RNP Granule Function. Neuron 88, 678–690 (2015).

20. Maharana, S. et al. RNA buffers the phase separation behavior of prion-like RNA binding proteins. Science 360, 918–921 (2018).

21. Qamar, S. et al. FUS Phase Separation Is Modulated by a Molecular Chaperone and Methylation of Arginine Cation-π Interactions. Cell 173, 720–734.e15 (2018).

22. St George-Hyslop, P. et al. The physiological and pathological biophysics of phase separation and gelation of RNA binding proteins in amyotrophic lateral sclerosis and fronto-temporal lobar degeneration. Brain Res. 1693, 11–23 (2018).

23. Wang, A. et al. A single N‐terminal phosphomimic disrupts TDP‐43 polymerization, phase separation, and RNA splicing. EMBO J. 37, e97452 (2018).

24. McGurk, L. et al. Poly(ADP-Ribose) Prevents Pathological Phase Separation of TDP-43 by Promoting Liquid Demixing and Stress Granule Localization. Mol. Cell 71, 703–717.e9 (2018).

25. Sabari, B. R. et al. Coactivator condensation at super-enhancers links phase separation and gene control. Science (80-.). 361, eaar3958 (2018).

26. Han, X. et al. Roles of the BRD4 short isoform in phase separation and active gene transcription. Nat. Struct. Mol. Biol. 27, 333–341 (2020).

27. Boija, A. et al. Transcription Factors Activate Genes through the Phase-Separation Capacity of Their Activation Domains. Cell 175, 1842–1855.e16 (2018).

28. Liao, Y. C. et al. RNA Granules Hitchhike on Lysosomes for Long-Distance Transport, Using Annexin A11 as a Molecular Tether. Cell 179, 147–164.e20 (2019).

29. Narayanan, T. & Kumar, A. Reentrant phase transitions in multicomponent liquid mixtures. Phys. Rep. 249, 135–218 (1994).

30. Banerjee, P. R., Milin, A. N., Moosa, M. M., Onuchic, P. L. & Deniz, A. A. Reentrant Phase Transition Drives Dynamic Substructure Formation in Ribonucleoprotein Droplets. Angew. Chemie Int. Ed. 56, 11354–11359 (2017).

31. Milin, A. N. & Deniz, A. A. Reentrant Phase Transitions and Non-Equilibrium Dynamics in Membraneless Organelles. Biochemistry 57, 2470–2477 (2018).

32. Choi, J. M., Dar, F. & Pappu, R. V. LASSI: A lattice model for simulating phase transitions of multivalent proteins. PLoS Comput. Biol. 15, e1007028 (2019).

33. Burke, K. A., Janke, A. M., Rhine, C. L. & Fawzi, N. L. Residue-by-Residue View of In Vitro FUS Granules that Bind the C-Terminal Domain of RNA Polymerase II. Mol. Cell 60, 231–241 (2015).

34. Loo, W. S. et al. Reentrant phase behavior and coexistence in asymmetric block copolymer electrolytes. Soft Matter 14, 2789–2795 (2018).

35. Zhang, F. et al. Reentrant condensation of proteins in solution induced by multivalent counterions. Phys. Rev. Lett. 101, 148101 (2008).

36. Zhang, F. et al. Universality of protein reentrant condensation in solution induced by multivalent metal ions. Proteins Struct. Funct. Bioinforma. 78, 3450–3457 (2010).

37. Zhang, F. et al. Reentrant condensation, liquid-liquid phase separation and crystallization in protein solutions induced by multivalent metal ions. Pure Appl. Chem. 86, 191–202 (2014).

38. Roosen-Runge, F., Heck, B. S., Zhang, F., Kohlbacher, O. & Schreiber, F. Interplay of pH and binding of multivalent metal ions: Charge inversion and reentrant condensation in protein solutions. J. Phys. Chem. B 117, 5777–5787 (2013).

39. Braun, M. K. et al. Reentrant Phase Behavior in Protein Solutions Induced by Multivalent Salts: Strong Effect of Anions Cl– Versus NO3–. J. Phys. Chem. B 122, 11978–11985 (2018).

40. Li, T., Ci, T., Chen, L., Yu, L. & Ding, J. Salt-induced reentrant hydrogel of poly(ethylene glycol)–poly(lactide-co-glycolide) block copolymers. Polym. Chem. 5, 979–991 (2014).

41. Mason, B. D., Zhang-van Enk, J., Zhang, L., Remmele, R. L. & Zhang, J. Liquid-Liquid Phase Separation of a Monoclonal Antibody and Nonmonotonic Influence of Hofmeister Anions. Biophys. J. 99, 3792–3800 (2010).

42. Dumetz, A. C., Chockla, A. M., Kaler, E. W. & Lenhoff, A. M. Protein Phase Behavior in Aqueous Solutions: Crystallization, Liquid-Liquid Phase Separation, Gels, and Aggregates. Biophys. J. 94, 570–583 (2008).

43. Taratuta, V. G., Holschbach, A., Thurston, G. M., Blankschtein, D. & Benedek, G. B. Liquid-liquid phase separation of aqueous lysozyme solutions: Effects of pH and salt identity. J. Phys. Chem. 94, 2140–2144 (1990).

44. Banani, S. F. et al. Compositional Control of Phase-Separated Cellular Bodies. Cell 166, 651–663 (2016).

45. Banjade, S. & Rosen, M. K. Phase transitions of multivalent proteins can promote clustering of membrane receptors. Elife 3, (2014).

46. Nott, T. J. et al. Phase Transition of a Disordered Nuage Protein Generates Environmentally Responsive Membraneless Organelles. Mol. Cell 57, 936–947 (2015).

47. Elbaum-Garfinkle, S. et al. The disordered P granule protein LAF-1 drives phase separation into droplets with tunable viscosity and dynamics. Proc. Natl. Acad. Sci. U. S. A. 112, 7189–7194 (2015).

48. Baldwin, R. L. How Hofmeister ion interactions affect protein stability. Biophys. J. 71, 2056–2063 (1996).

49. Pegram, L. M. et al. Why Hofmeister effects of many salts favor protein folding but not DNA helix formation. Proc. Natl. Acad. Sci. U. S. A. 107, 7716–7721 (2010).

50. Kohn, W. D., Kay, C. M. & Hodges, R. S. Salt effects on protein stability: Two-stranded α-helical coiled-coils containing inter- or intrahelical ion pairs. J. Mol. Biol. 267, 1039–1052 (1997).

51. Beauchamp, D. L. & Khajehpour, M. Studying salt effects on protein stability using ribonuclease t1 as a model system. Biophys. Chem. 161, 29–38 (2012).

52. Duong-Ly, K. C. & Gabelli, S. B. Salting out of Proteins Using Ammonium Sulfate Precipitation. Methods Enzymol. 85–94 (2014). doi:10.1016/B978-0-12-420119-4.00007-0

53. Arakawa, T. & Timasheff, S. N. Mechanism of Protein Salting In and Salting Out by Divalent Cation Salts: Balance between Hydration and Salt Binding. Biochemistry 23, 5912–5923 (1984).

54. Curtis, R. A., Prausnitz, J. M. & Blanch, H. W. Protein‐protein and protein‐salt interactions in aqueous protein solutions containing concentrated electrolytes. Biotechnol. Bioeng. 57, 11–21 (1998).

55. Dumetz, A. C., Snellinger-O’Brien, A. M., Kaler, E. W. & Lenhoff, A. M. Patterns of protein-protein interactions in salt solutions and implications for protein crystallization. Protein Sci. 16, 1867–1877 (2007).

56. Leirmo, S., Harrison, C., Cayley, D. S., Record, M. T. & Burgess, R. R. Replacement of Potassium Chloride by Potassium Glutamate Dramatically Enhances Protein-DNA Interactions in Vitro. Biochemistry 26, 2095–2101 (1987).

57. Murdoch, F. E., Grunwald, K. A. A. & Gorski, J. Marked Effects of Salt on Estrogen Receptor Binding to DNA: Biologically Relevant Discrimination between DNA Sequences. Biochemistry 30, 10838–10844 (1991).

58. Xiao, B., Johnson, R. C. & Marko, J. F. Modulation of HU-DNA interactions by salt concentration and applied force. Nucleic Acids Res. 38, 6176–85 (2010).

59. Li, J. et al. BET bromodomain inhibition promotes neurogenesis while inhibiting gliogenesis in neural progenitor cells. Stem Cell Res. 17, 212–221 (2016).

60. Ferri, A. L. M. et al. Sox2 deficiency causes neurodegeneration and impaired neurogenesis in the adult mouse brain. Development 131, 3805–3819 (2004).

61. Monahan, Z. et al. Phosphorylation of the FUS low‐complexity domain disrupts phase separation, aggregation, and toxicity. EMBO J. 36, 2951–2967 (2017).

62. Murthy, A. C. et al. Molecular interactions underlying liquid−liquid phase separation of the FUS low-complexity domain. Nat. Struct. Mol. Biol. 26, 637–648 (2019).

63. Kato, M. et al. Cell-free formation of RNA granules: Low complexity sequence domains form dynamic fibers within hydrogels. Cell 149, 753–767 (2012).

64. Patel, A. et al. ATP as a biological hydrotrope. Science 356, 753–756 (2017).

65. Kang, J., Lim, L. & Song, J. ATP enhances at low concentrations but dissolves at high concentrations liquid-liquid phase separation (LLPS) of ALS/FTD-causing FUS. Biochem. Biophys. Res. Commun. 504, 545–551 (2018).

66. Nishi, H., Hashimoto, K. & Panchenko, A. R. Phosphorylation in protein-protein binding: Effect on stability and function. Structure 19, 1807–1815 (2011).

67. Bannister, A. J. & Kouzarides, T. Regulation of chromatin by histone modifications. Cell Res. 21, 381–395 (2011).

68. Berne, B. J., Weeks, J. D. & Zhou, R. Dewetting and Hydrophobic Interaction in Physical and Biological Systems. Annu. Rev. Phys. Chem. 60, 85–103 (2009).

69. Dahal, Y. R. & Schmit, J. D. Ion Specificity and Nonmonotonic Protein Solubility from Salt Entropy. Biophys. J. 114, 76–87 (2018).

70. Kroschwald, S., Maharana, S. & Simon, A. Hexanediol: a chemical probe to investigate the material properties of membrane-less compartments. Matters (2017). doi:10.19185/matters.201702000010

71. Boeynaems, S. et al. Phase Separation of C9orf72 Dipeptide Repeats Perturbs Stress Granule Dynamics. Mol. Cell 65, 1044–1055.e5 (2017).

72. Boeynaems, S. et al. Spontaneous driving forces give rise to protein-RNA condensates with coexisting phases and complex material properties. Proc. Natl. Acad. Sci. 116, 7889–7898 (2019).

73. Hofmeister, F. Zur Lehre von der Wirkung der Salze. Arch. für Exp. Pathol. und Pharmakologie 24, 247–260 (1888).

74. Cacace, M. G., Landau, E. M. & Ramsden, J. J. The Hofmeister series: salt and solvent effects on interfacial phenomena. Q. Rev. Biophys. 30, 241–77 (1997).

75. Hyde, A. M. et al. General Principles and Strategies for Salting-Out Informed by the Hofmeister Series. Org. Process Res. Dev. 21, 1355–1370 (2017).

76. Mazzini, V. & Craig, V. S. J. What is the fundamental ion-specific series for anions and cations? Ion specificity in standard partial molar volumes of electrolytes and electrostriction in water and non-aqueous solvents. Chem. Sci. 8, 7052–7065 (2017).

77. Best, R. B., Zheng, W. & Mittal, J. Balanced Protein–Water Interactions Improve Properties of Disordered Proteins and Non-Specific Protein Association. J. Chem. Theory Comput. 10, 5113–5124 (2014).

78. Benavides, A. L., Aragones, J. L. & Vega, C. Consensus on the solubility of NaCl in water from computer simulations using the chemical potential route. J. Chem. Phys. 144, 124504 (2016).

79. Minoux, H. & Chipot, C. Cation-π interactions in proteins: Can simple models provide an accurate description? J. Am. Chem. Soc. 121, 10366–10372 (1999).

80. Khan, H. M. et al. Improving the Force Field Description of Tyrosine–Choline Cation−π Interactions: QM Investigation of Phenol–N(Me)4+ Interactions. J. Chem. Theory Comput. 12, 5585–5595 (2016).

81. Caldwell, J. W. & Kollman, P. A. Cation-.pi. Interactions: Nonadditive Effects Are Critical in Their Accurate Representation. J. Am. Chem. Soc. 117, 4177–4178 (1995).

82. Demerdash, O., Mao, Y., Liu, T., Head-Gordon, M. & Head-Gordon, T. Assessing many-body contributions to intermolecular interactions of the AMOEBA force field using energy decomposition analysis of electronic structure calculations. J. Chem. Phys. 147, 161721 (2017).

83. Cubero, E., Luque, F. J. & Orozco, M. Is polarization important in cation-π interactions? Proc. Natl. Acad. Sci. U. S. A. 95, 5976–5980 (1998).

84. Shi, Z., Olson, C. A. & Kallenbach, N. R. Cation-π interaction in model α-helical peptides. J. Am. Chem. Soc. 124, 3284–3291 (2002).

85. Shi, Z., Olson, C. A., Bell, A. J. & Kallenbach, N. R. Stabilization of α-helix structure by polar side-chain interactions: Complex salt bridges, cation-π interactions, and C-H···O H-bonds. Biopolymers - Peptide Science Section 60, 366–380 (2001).

86. Xie, N.-Z., Du, Q.-S., Li, J.-X. & Huang, R.-B. Exploring Strong Interactions in Proteins with Quantum Chemistry and Examples of Their Applications in Drug Design. PLoS One 10, e0137113 (2015).

87. Dyson, H. J., Wright, P. E. & Scheraga, H. A. The role of hydrophobic interactions in initiation and propagation of protein folding. Proc. Natl. Acad. Sci. U. S. A. 103, 13057–13061 (2006).

88. Andrew, C. D. et al. Stabilizing interactions between aromatic and basic side chains in α-helical peptides and proteins. Tyrosine effects on helix circular dichroism. J. Am. Chem. Soc. 124, 12706–12714 (2002).

89. Vernon, R. M. C. et al. Pi-Pi contacts are an overlooked protein feature relevant to phase separation. Elife 7, (2018).

90. Fisher, R. S. & Elbaum-Garfinkle, S. Tunable multiphase dynamics of arginine and lysine liquid condensates. Nat. Commun. 11, 4628 (2020).

91. Tesei, G. et al. Self-association of a highly charged arginine-rich cell-penetrating peptide. Proc. Natl. Acad. Sci. 114, 11428–11433 (2017).

92. Dignon, G. L., Zheng, W., Best, R. B., Kim, Y. C. & Mittal, J. Relation between single-molecule properties and phase behavior of intrinsically disordered proteins. Proc. Natl. Acad. Sci. 115, 9929–9934 (2018).

93. Dignon, G. L., Zheng, W., Kim, Y. C. & Mittal, J. Temperature-Controlled Liquid-Liquid Phase Separation of Disordered Proteins. ACS Cent. Sci. 5, 821–830 (2019).

94. Kang, J., Lim, L., Lu, Y. & Song, J. A unified mechanism for LLPS of ALS/FTLD-causing FUS as well as its modulation by ATP and oligonucleic acids. PLoS Biol. 17, e3000327 (2019).

95. Adame-Arana, O., Weber, C. A., Zaburdaev, V., Prost, J. & Jülicher, F. Liquid phase separation controlled by pH. (2019).

96. Ruff, K. M., Roberts, S., Chilkoti, A. & Pappu, R. V. Advances in Understanding Stimulus-Responsive Phase Behavior of Intrinsically Disordered Protein Polymers. J. Mol. Biol. 430, 4619–4635 (2018).

97. Quiroz, F. G. & Chilkoti, A. Sequence heuristics to encode phase behaviour in intrinsically disordered protein polymers. Nat. Mater. 14, 1164–1171 (2015).

98. Cinar, H. et al. Temperature, Hydrostatic Pressure, and Osmolyte Effects on Liquid– Liquid Phase Separation in Protein Condensates: Physical Chemistry and Biological Implications. Chem. – A Eur. J. 25, 13049–13069 (2019).

99. Cinar, H., Cinar, S., Chan, H. S. & Winter, R. Pressure-Induced Dissolution and Reentrant Formation of Condensed, Liquid-Liquid Phase-Separated Elastomeric α-Elastin. Chem. - A Eur. J. 24, 8286–8291 (2018).

100. Wheeler, R. J. et al. Small molecules for modulating protein driven liquid-liquid phase separation in treating neurodegenerative disease. bioRxiv 721001 (2019). doi:10.1101/721001

101. Hughes, M. P. et al. Atomic structures of low-complexity protein segments reveal kinked β sheets that assemble networks. Science 359, 698–701 (2018).

