## Supplementary Information for "Reentrant liquid condensate phase of proteins is stabilized by hydrophobic and non-ionic interactions"

**The Supplementary Information includes:**

Methods (Materials, Protein production, Sample preparation and generation of phase diagrams, Fluorescence imaging, Simulation Methods)
Supplementary Information for atomistic PMF calculations (Tables S1 and S2)
Supplementary Information for condensate densities (Table S3)
Supplementary Figures (Figures S1, S2, S3, S4)
Supplementary References

### Methods

#### Materials

All reagents and chemicals were purchased with the highest purity available. PolyU RNA with a molecular weight range from 800–1,000 kDa was purchased from Sigma Aldrich as lyophilized powder and dissolved into a stock of 5 mg/mL in 50 mM Tris-HCl (pH 7.2) before use. ATP was obtained from Fisher Scientific and a 50 mM stock solution was prepared in 50 mM Tris-HCl (pH 7.2). 1,6-Hexanediol was purchased from Santa Cruz Biotechnology Inc. and a 40% (*w*/*v*) stock solution was prepared in 50 mM Tris-HCl (pH 7.2). KCl was from Fisher Scientific and all the chloride salts used in the Hofmeister series experiments were from Sigma Aldrich. The $PR_{25}$ peptide containing 25 proline–arginine repeats was obtained from GenScript. N-terminally labelled $PR_{25}$ was obtained by reacting the peptide with amine-reactive AlexaFluor546 (Sigma Aldrich).

#### Protein production

**FUS, FUS G156E, TDP-43, and Annexin A11.** FUS and FUS G156E were produced as C-terminal EGFP fusion proteins from Sf9 insect cells as previously described[1] and stored at a concentration of 75 μM in 50 mM Tris-HCl (pH 7.4), 500 mM KCl, 1 mM DTT, 5% glycerol. TDP-43 was similarly produced as a C-terminal GFP fusion protein as previously described[2] and stored at 120 μM in the same buffer as the FUS proteins. Annexin A11 was expressed and purified from insect Sf9 cells using standard procedures as previously described, labelled with Alexa647 and stored at a concentration of 150 μM in 20 mM HEPES (pH 7.0), 225 mM NaCl.[3]

**Sox2 and Brd4.** Proteins were expressed in Sf9 insect cells for 72 h using the baculovirus system.[4] Brd4 and Sox2 were produced as N-/C-terminal monoGFP fusion proteins, respectively, as described below in detail. After purification, proteins were aliquoted, snap-frozen in liquid nitrogen and stored at –80°C until usage.

*Sox2.* Cells expressing $His_6$-MBP-Sox2-GFP were resuspended in Sox2 buffer (50 mM Bis-Tris-Propane pH 7.5, 500 mM KCl, 5% glycerol, 1 mM DTT) supplemented with EDTA-free protease inhibitor cocktail set III (Calbiochem) and 0.25 U/mL Benzonase (in-house) and lysed by sonication. The lysate was clarified by centrifugation for 2.5 h at 13,000 *g* and 4°C. The supernatant was applied to an Econo-Pac gravity columns (Bio-Rad) filled with amylose resin (NEB). After washing the beads with 3 column volumes of Sox2 buffer, the protein was eluted with Sox2 buffer supplemented with 50 mM maltose. The $His_6$-MBP moiety was removed by incubation with 140 U 3C preScission protease (in-house) on a rotator for 3 h at room temperature. After concentrating of the protein to about 2.5 mL using a 50,000 MWCO Amicon Ultra concentrator (Millipore), the protein was diluted again with a solution containing 50 mM Bis-Tris-Propane pH 7.5, 5% glycerol and 1 mM DTT to a final KCl concentration of 100 mM. Next, cation exchange was performed using a HiTrap SP HP column (GE Healthcare) and 50 mM Bis-Tris-Propane pH 7.4, 5% glycerol and 1 mM DTT as a buffer with a KCl gradient ranging from 100 mM to 1 M. Fractions containing Sox2-GFP were pooled and subjected to size exclusion chromatography using a Superdex 200 Increase 10/300 GL column (GE Healthcare) and Sox2. As before, protein-containing fractions were pooled and concentrated with a 30,000 MWCO Amicon Ultra concentrator (Millipore).

*Brd4.* Cells expressing $His_6$-MBP-GFP-Brd4 were resuspended in Brd4 buffer (50 mM Tris pH 7.5, 500 mM KCl, 5 mM EDTA, 5% glycerol) supplemented with EDTA-free protease inhibitor cocktail set III (Calbiochem) and 0.25 U/mL Benzonase (in-house) and lysed by sonication. The lysate was clarified by centrifugation for 1 h at 13,000 *g* and 4°C. The supernatant was applied to an Econo-Pac gravity columns (Bio-Rad) filled with amylose resin (NEB). After washing the beads with 3 column volumes of Brd4 buffer, the protein was eluted with Brd4 buffer supplemented with 50 mM maltose. The $His_6$-MBP moiety was removed by incubation with 70 U 3C preScission protease (in-house) on a rotator for 2 h at room temperature. After concentrating of the protein using a 30,000 MWCO Amicon Ultra concentrator (Millipore), the protein subjected to size exclusion chromatography using a Superdex 200 Increase 10/300 GL column (GE Healthcare) and Brd4 storage buffer (50 mM Tris pH 7.5, 500 mM KCl, 2 mM EDTA, 5% glycerol, 1 mM DTT). As before, protein-containing fractions were pooled and concentrated with a 30,000 MWCO Amicon Ultra concentrator (Millipore).

**Sample preparation and generation of phase diagrams**

To induce phase separation of proteins, appropriate amounts of salt, water, and additives, as indicated, were added to the protein stock solutions and mixed by pipetting. In all cases the buffer contained 50 mM Tris-HCl (pH 7.2), except for Brd4 and Sox2, which contained 5 mM Tris (pH 7.5), 0.2 mM EDTA, 0.5% glycerol and 5 mM Bis-Tris-Propane (pH 7.5), 0.5% glycerol, respectively. Phase separated samples were prepared in tubes and imaged within 1–5 minutes to limit any ageing effects; Brd4 and Sox2 were imaged after an incubation period of 10–20 minutes.

Phase diagrams of FUS and $PR_{25}$ were generated by mixing protein/peptide stocks with the respective salt solutions in 50 mM Tris-HCl (pH 7.2), as described above. The resulting sample solutions were then imaged immediately after preparation. Phase diagrams were constructed by systematically screening through conditions and assessing conditions in which a dense phase or a well-mixed state was detected. Regions close to the phase boundary were mapped out and each point on the phase diagram was tested at least twice.

**Fluorescence imaging**

Imaging of FUS, FUS G156E, TDP-43, A11 and $PR_{25}$ samples was performed on an inverted fluorescence microscope (Zeiss AxioObserver D1) equipped with a high-sensitivity camera (Evolve 512 EMCCD, Photometrics) by placing an aliquot of the sample (3 μL) on a microscope slide mounted on the microscope stage. FUS, FUS G156E, TDP-43, and $PR_{25}$ samples were imaged using a Zeiss A-Plan 20x/0.30 NA air objective; A11 samples were imaged on a Zeiss A-Plan 100x/1.25 NA oil-immersion objective. In experiments with FUS proteins and TDP-43, an appropriate filter set for GFP detection was used (49002, Chroma Technology). Similarly, labelled $PR_{25}$ was detected with a filter set for AlexaFluor546 fluorescence detection (49004, Chroma Technology). A11 was imaged with a filter set for AlexaFluor647 fluorescence detection (49009, Chroma Technology). Imaging of Brd4 and Sox2 was performed on a Nikon-Andor Eclipse Ti inverted spinning disc confocal microscope equipped with an Andor iXON 897 EMCCD camera and a 60x/1.2 NA water-immersion objective (Nikon). Brd4 and Sox2 samples were transferred to glass slides that were pre-prepared by first cutting a strip of double-sided tape (Scotch) into 1x1 cm squares which were then stuck onto the glass slides. 2 µL of the sample was pipetted onto each slide and a PEGylated cover slip was used to seal the chamber in the distance of the tape. For experiments involving dissolution induced by additives, components were mixed 3:1 with 3 μL of phase separated protein at the specified salt concentration with 1 μL of the additional component (*i.e.*, 1,6-hexanediol, PolyU RNA, ATP). The final concentrations of protein and additional components are stated in each figure.

**Simulation methods**

**PMF calculations.** PMF calculations were carried out using the GROMACS simulation package (version 2019.3).[5] Amino acids were modelled using the AMBERff03ws force field.[6] Since, we are interested in probing interaction potentials at very high salt concentrations (up to 3 M NaCl), it is very important that the solvent and ion model parameters employed are well-fitted to fairly reproduce ion solubilities in water at 298 K (*i.e.*, an absence of unphysical salt crystallization). The JC-SPC/E-ion/TIP4P/2005 force field has been optimized for that purpose, and so it was used in this work.[7] The N- and C-terminal ends of each amino acid were capped with acetyl and an N-methyl capping groups, respectively. Pairs of amino acids were oriented with their sidechains facing each other, based on the most common arrangements observed in protein structures. Dimers were immersed in a cubic box containing TIP4P/2005 water molecules (ca. 1400–3400 molecules) with a minimum distance of 1.0 nm between the dimer and the edge of the box. When necessary, some water molecules were replaced by $Na^+$ and/or $Cl^-$ ions to produce neutral systems. Energy minimizations (force tolerance = 500 kJ mol$^{-1}$ nm$^{-1}$) were performed for the neutralized systems, with positional restraints of 20,000 kJ mol$^{-1}$ nm$^{-2}$ applied in each dimension to all amino acid heavy-atoms. $Na^+$ and $Cl^-$ ions were then added to yield the desired salt concentrations (0 M, 0.15 M, 1.5 M, 3 M).

For production runs, positional restraints of 1000 kJ mol$^{-1}$ nm$^{-2}$ (in the directions perpendicular to the pulling direction) were used to constrain amino acid heavy-atoms. The center of mass (COM) distance between amino acid pairs was restrained with a harmonic umbrella potential (pulling force constant = 6000 kJ mol$^{-1}$ nm$^{-2}$). Bonds with hydrogens were constrained using the LINCS algorithm, permitting an integration timestep of 2 fs. Periodic boundary conditions (PBC) were used during MD simulations and electrostatics were computed using Particle-Mesh Ewald summations[8] with a Coulomb cutoff of 0.9 nm. For each concentration, approximately 30 windows, spaced at 0.05 nm from 0.1 to 1.6 nm, were used per amino acid pair. Each window was simulated for 10 ns. Three independent simulations were conducted for each umbrella sampling window (*i.e.*, an aggregate simulation time of 30 ns per window). Umbrella sampling simulations were analyzed using WHAM,[9] as implemented in GROMACS. The first 1000 ps of simulations were used for equilibration and were not included in the WHAM analysis. Error analysis was performed using the Bayesian bootstrap method, as described by Hub and coworkers.[10]

**Cation–π charge refitting.** To study the impact of salt on cation–π interactions, which are not well captured by standard atomistic force fields[11] because they involve the polarization of the π-electron cloud[12] of an aromatic side chain (Tyr, Phe, Trp) due to the cationic side chain (Lys, Arg), we refitted the charges on the Tyr and Phe sidechains when bound to Arg or Lys. We first performed constrained (*i.e.*, backbone and capping group heavy-atoms were frozen) geometry optimizations of each dimer (Arg–Tyr, Arg–Phe, Lys–Phe) at the MP2/6-31G(d) level of theory using the Gaussian 09 program.[13] The electrostatic surface potential (ESP) was then computed for each optimized pair. Finally, the sidechain charges of Tyr and Phe were refitted based on the quantum mechanical EPSs using the RESP program in Amber (maintaining the charge symmetry in the rings).[14]

**Coarse-grained protein model.** We have implemented a reparametrized version (see '*Experimental validation and refinement of coarse-grained model'* below) of the sequence-dependent coarse-grained model of the Mittal group, originally developed to capture qualitatively the sequence-dependent phase behavior of proteins that undergo LLPS at physiological salt conditions (~100 mM NaCl).[15] The model implements a resolution of one bead per amino acid and a sequence-dependent set of parameters derived top-down to approximate the single-molecule experimental radius of gyration of a wide-range of intrinsically disordered proteins. Intrinsically disordered protein regions are treated as flexible polymers and globular regions as rigid bodies. Inter-residue bonds within the disordered domains are described using harmonic springs. Longrange electrostatics are modelled using a Coulombic term with Debye–Hückel electrostatic screening. Nonbonded pairwise interactions are modelled using a knowledge-based potential termed hydrophobicity scale (HPS) model that is based on one of the hydrophobicity scales for amino acids available.[16] For the globular protein domains, a 30% scaled down set of the HPS parameters was used to account for 'buried' amino acids. Because the model distinguishes between disordered and globular protein regions and maintains the secondary structure of globular regions, it requires an initial atomistic model for the proteins; these are described below.

**Initial atomistic models for coarse-grained simulations.** We simulated the phase behavior of the full length FUS protein (Uniprot code: K7DPS7, 526 residues, 24 proteins), the prion like domain (PLD) of FUS for validation only (residues: 1–163, 100 proteins), and a reduced version of the $PR_{25}$ protein (13 Arg and 12 Pro residues alternately positioned, 400 proteins). Since the structure of full length FUS has not been resolved, we developed an atomistic model by fusing the intrinsically disordered regions with the resolved structural domains (residues from 285–371 (PDB code: 2LCW) and from 422–453 (PDB code: 6G99)). An initial intrinsically disordered model for $PR_{25}$ was developed in VMD.[17]

**Coarse-grained simulation methods.** To evaluate the formation of liquid condensates in the different systems, we performed direct coexistence simulations[18–20] at constant volume and temperature. The direct coexistence method simulates the condensate and diluted phases in the same box separated by an interface. The initial simulation box was prepared by running simulations at constant temperature and a pressure of 1 bar, using the Berendsen barostat, and then enlarging the simulation box in one direction ~4 times. The simulation temperatures were chosen to be just below the correspondent critical temperatures for each system: 400 K for full length FUS and 200 K for $PR_{25}$. For the production runs, each system was simulated for ~2.5 μs, using a Langevin thermostat with relaxation time of 5 ps and a time step of 10 fs.[21] The LAMMPS software MD package was used to carry out all the coarse-grained simulations.[22]

**Experimental validation and refinement of coarse-grained model.** To verify that the HPS model captures qualitatively the experimental phase behavior of FUS at low salt, we first used it to compute the phase diagrams of the PLD of FUS and the full FUS protein. While experiments demonstrate a higher LLPS propensity of the full FUS protein than of the PLD,[23,24] our simulations with the original parametrization found a ~20% higher critical point for the PLD of FUS versus full FUS. The PLD is almost completely devoid of charged residues but rich in Tyr, and its LLPS has been shown experimentally to be stabilized by hydrophobic forces.[25] In contrast, full FUS contains three Arg-rich motifs, and experiments demonstrate that its LLPS at low salt is dependent on Arg–Tyr cation–π interactions,[26] and electrostatic screening inhibits LLPS (as demonstrated experimentally in this work). These findings suggested that enhanced cation–π and electrostatic interactions were required to achieve qualitative agreement with the FUS experimental behavior. When we included an additional term in the potential energy to increase the strength of cation–π interactions at low salt, as recently proposed,[27] we qualitatively recover a higher critical temperature for full FUS versus its PLD. Our PMFs indicate that when transitioning from low to moderate salt, electrostatic interactions diminish significantly, while hydrophobic interactions remain strong, giving rise to reentrant phase behavior. However, when we tested the HPS+cation–π enhanced model in the extreme scenario of no electrostatic contribution to the potential energy and constant hydrophobicity, we observed no statistically significant difference in the critical temperature of FUS (with respect to the normal HPS+cation–π enhanced model); suggesting that this combination of parameters now underestimates the relative electrostatic contribution to the potential energy for FUS. We thus investigated the modulation of the phase diagram of full FUS in the HPS+cation–π enhanced model versus the relative electrostatic contribution to the potential energy by multiplying the Coulomb interaction by a parameter $\chi$ = 0, 1, 2 and 4. We found qualitatively similar phase diagrams for $\chi$ = 0, 1 and 2, and a 4% increase in the critical temperature for FUS with $\chi$ = 4; suggesting that in the HPS+cation–π enhanced model with $\chi$=0, 1 and 2 electrostatics still do not play a significant role in the phase behavior of FUS. Hence, to mimic low salt conditions, we increased the electrostatic contribution to the potential energy by a factor of four ($\chi$ = 4) and used this as our reference model. With this reparameterization, we recomputed the phase diagrams of full FUS and its PLD at low salt, and obtained a convincingly higher critical point for the full FUS (~15%) with respect to that of the PLD, in qualitative agreement with experimental observations.[23,24] Based on the salt-dependent trends from our PMFs, we approximate the moderate salt regime (1.5–3 M NaCl) by scaling down the strength of electrostatic interactions with respect to our reference model (*i.e.*, we set $\chi$ = 2), and the high salt regime (>3 M NaCl) by setting $\chi$ = 1 and increasing the hydrophobic contribution by 10–30%.

**Estimation of contact frequencies from coarse-grained simulations.** Average number of protein contacts within phase-separated condensates were calculated using the MDAnalysis Python library.[28,29] Amino acids in two different proteins are in contact if they are within a cut-off distance of 0.65 nm of each other. Using this criterion, we estimated the frequency of contacts between the different domains of FUS (see Figure S4).

**Supplementary Information for atomistic PMF calculations**

**Table S1:** Refitted charges for Tyr in Arg–Tyr dimer and Phe in Arg–Phe and Lys–Phe dimers. All dimers are in the parallel geometry (as shown in Figure 6g of main text and in Figure S2 below). The original and refitted force field parameters are summarized below.

| Atom in Tyr | Original charges | Refitted Tyr charges in Arg–Tyr | Atom in Phe | Original charges | Refitted Phe charges in Arg–Phe | Refitted Phe charges in Lys–Phe |
|---|---|---|---|---|---|---|
| **CB** | -0.051853 | 0.516775 | **CB** | -0.09872 | 0.755677 | 0.946836 |
| **HB1** | 0.019145 | -0.154966 | **HB1** | 0.060989 | -0.209079 | -0.25544 |
| **HB2** | 0.019145 | -0.154966 | **HB2** | 0.060989 | -0.209079 | -0.25544 |
| **CG** | 0.112601 | -0.220971 | **CG** | 0.021313 | -0.203187 | -0.387676 |
| **CD1** | -0.183461 | -0.125181 | **CD1** | -0.083109 | -0.273037 | -0.25241 |
| **HD1** | 0.132715 | 0.130310 | **HD1** | 0.098466 | 0.128345 | 0.221075 |
| **CE1** | -0.181823 | -0.299333 | **CE1** | -0.156974 | 0.027607 | -0.211827 |
| **HE1** | 0.137303 | 0.202092 | **HE1** | 0.123731 | 0.095427 | 0.167398 |
| **CZ** | 0.206277 | 0.337235 | **CZ** | -0.099824 | -0.22782 | 0.052773 |
| **OH** | -0.421233 | -0.492286 | **HZ** | 0.114679 | 0.160458 | 0.074129 |
| **HH** | 0.329691 | 0.376644 | **CE2** | -0.156974 | 0.027607 | -0.211827 |
| **CE2** | -0.181823 | -0.299333 | **HE2** | 0.123731 | 0.095427 | 0.167398 |
| **HE2** | 0.137303 | 0.202092 | **CD2** | -0.083109 | -0.273037 | -0.25241 |
| **CD2** | -0.183461 | -0.125181 | **HD2** | 0.098466 | 0.128345 | 0.221075 |
| **HD2** | 0.132715 | 0.130310 | | | | |

**Table S2:** Refitted charges for Tyr in Arg–Tyr and Phe in Lys–Phe for amino acid pairs in t-shaped geometries (see Figure S3 below). The original and refitted force field parameters are summarized below.

| Atom in Tyr | Original charges | Refitted Tyr charges in Arg–Tyr | Atom in Phe | Original charges | Refitted Phe charges in Lys–Phe |
|---|---|---|---|---|---|
| **CB** | -0.051853 | 0.662808 | **CB** | -0.09872 | 0.963787 |
| **HB1** | 0.019145 | -0.177211 | **HB1** | 0.060989 | -0.28587 |
| **HB2** | 0.019145 | -0.177211 | **HB2** | 0.060989 | -0.28587 |
| **CG** | 0.112601 | -0.612723 | **CG** | 0.021313 | -0.323652 |
| **CD1** | -0.183461 | 0.185225 | **CD1** | -0.083109 | -0.334344 |
| **HD1** | 0.132715 | 0.14026 | **HD1** | 0.098466 | 0.214518 |
| **CE1** | -0.181823 | -0.626374 | **CE1** | -0.156974 | -0.001983 |
| **HE1** | 0.137303 | 0.25684 | **HE1** | 0.123731 | 0.123162 |
| **CZ** | 0.206277 | 0.824476 | **CZ** | -0.099824 | -0.187325 |
| **OH** | -0.421233 | -0.881568 | **HZ** | 0.114679 | 0.139878 |
| **HH** | 0.329691 | 0.472768 | **CE2** | -0.156974 | -0.001983 |
| **CE2** | -0.181823 | -0.626374 | **HE2** | 0.123731 | 0.123162 |
| **HE2** | 0.137303 | 0.25684 | **CD2** | -0.083109 | -0.334344 |
| **CD2** | -0.183461 | 0.185225 | **HD2** | 0.098466 | 0.214518 |
| **HD2** | 0.132715 | 0.14026 | | | |

### Supplementary Information for condensate densities

**Table S3:** Densities of FUS condensates computed using coarse-grained model parameters for different salt regimes. Statistical uncertainties are given in parentheses. n.a., not applicable.

| Salt regime | Density of condensate (g/cm$^3$) |
|---|---|
| Low salt | 0.33 (7%) |
| Medium salt | n.a. |
| High salt | 0.50 (7%)* |

* Sensibly higher density, which means that we have crossed the coexistence line; thus, an increase in hydrophobicity of 5% (versus 10%, as described in the main text) would have been sufficient to reenter the condensed phase.

### Supplementary Figures

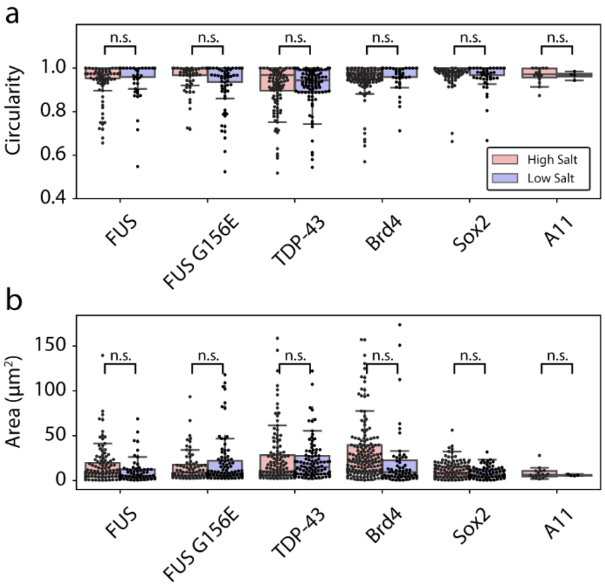


**Figure S1. Analysis of condensate circularity and condensate area in the low and high salt regimes for FUS, FUS G156E, TDP-43, Brd4, Sox2, and A11. (a)** Analysis of condensate circularities. For all proteins, median circularities were >0.95. For each protein, at high salt and low salt, respectively, the median circularities were: FUS: 1.00, 1.00; FUS G156E: 1.00, 1.00; TDP-43: 0.97, 0.94; Brd4: 0.98, 1.00; Sox2: 1.00, 1.00; A11: 0.97, 0.97. Statistical analysis was performed using a two-sided *t*-test. *p*-values were found to be FUS: $p > 0.7518$; FUS G156E: $p > 0.0948$; TDP-43: $p > 0.6801$; Brd4: $p > 0.3082$; Sox2: $p > 0.1503$; A11: $p > 0.9973$. No significant difference in circularity was found between high- and low-salt condensates (n.s., not significant). **(b)** Analysis of condensate area. For each protein, at high salt and low salt, respectively, the median areas (reported in $\mu m^2$) were: FUS: 7.60, 2.95; FUS G156E: 4.60, 3.86; TDP-43: 6.70, 7.83; Brd2: 21.22, 9.43; Sox2: 8.91, 6.81; A11: 6.61, 5.87. Statistical analysis between the areas was performed using a two-sided *t*-test. *p*-values were found to be FUS: $p > 0.0645$; FUS G156E: $p > 0.2440$; TDP-43: $p > 0.3809$; Brd4: $p > 0.1710$; Sox2: $p > 0.0935$; A11: $p > 0.5040$. No significant difference in area (n.s., not significant) was found between high- and low-salt condensates. Overall, the dataset confirms the morphological similarities between condensates in the high- and low-salt regimes. Note: In each of the images the pixel resolution size is near the diffraction limit, due to the high threshold value used as a cut-off condition; hence, only condensates above the diffraction limit were used in accessing condensates circularity and size. Circularity analysis and area analysis was performed on subsections of the images shown in Fig. 2 (see main text). Circularity was calculated using the formula: $\text{circ} = 4\pi\left(\frac{\text{area}}{\text{perimeter}}\right)^2$. Image analysis was done in Fiji/ImageJ and statistical analysis was conducted in Python. $n_{\text{FUS, low salt}} = 60$, $n_{\text{FUS, high salt}} = 119$, $n_{\text{FUS G156E, low salt}} = 95$; $n_{\text{FUS G156E, high salt}} = 75$; $n_{\text{TDP-43, low salt}} = 100$; $n_{\text{TDP-43, high salt}} = 130$; $n_{\text{Brd4, low salt}} = 57$; $n_{\text{Brd4, high salt}} = 162$; $n_{\text{Sox2, low salt}} = 75$; $n_{\text{Sox2, high salt}} = 139$; $n_{\text{A11, low salt}} = 4$; $n_{\text{A11, high salt}} = 16$. In both plots, boxes extend from the 25th to 75th percentiles, with a line at the median. Whiskers span 1.5× the interquartile range. Source data are provided as a Source Data file.

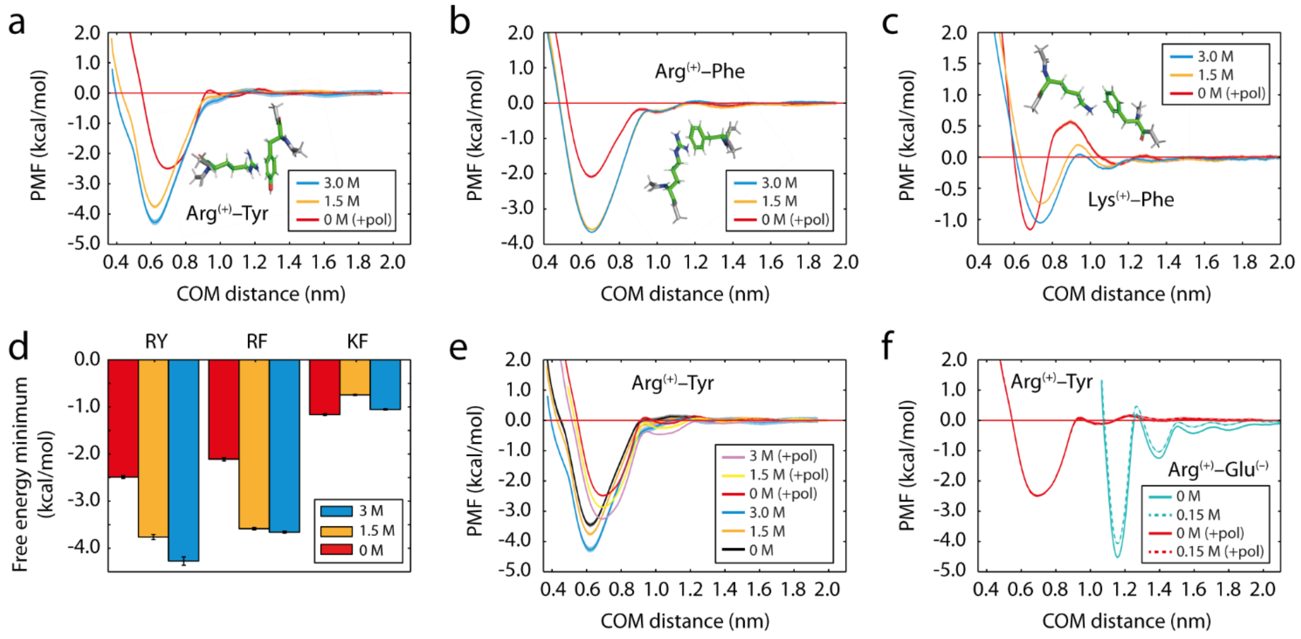


**Figure S2. Effect of salt on the potential of mean force (PMF) between basic and aromatic amino acid pairs in explicit solvent and NaCl ions as a function of the center-of-mass (COM) distance.** In the plots, +pol denotes refitted Tyr/Phe parameters were employed (as described above and summarized in Table S1). **(a)** Arg–Tyr (as in main text). **(b)** Arg–Phe. **(c)** Lys–Phe. Error bars are shown as bands and represent the standard deviations obtained by bootstrapping the results from three independent simulations. **(d)** Variation in the free energy minimum (obtained from the profiles in a–c**,** with associated error bars) with salt. One-letter amino acid codes are used to identify each pair interaction. **(e)** Comparison of Arg–Tyr interaction strength computed using the original force field Tyr parameters with those calculated using the modified Tyr parameters (i.e., obtained from the charge refitting procedure). **(f)** Arg–Tyr versus Arg–Glu interaction at low salt concentrations; plots reveal differences in sensitivity to electrostatic screening and variations in the interaction ranges.

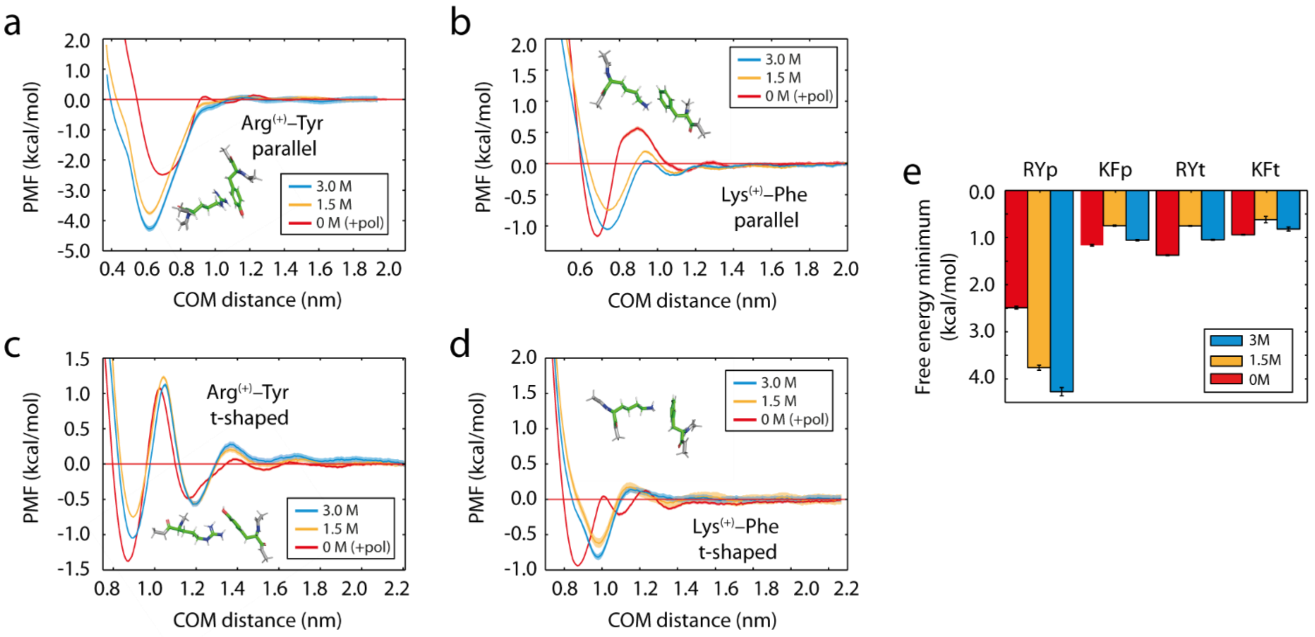


**Figure S3. Effects of geometry (parallel versus t-shaped) on the potential of mean force (PMF) between basic and aromatic amino acid pairs, computed at different salt concentrations, as a function of the center-of-mass (COM) distance.** In the plots, +pol denotes refitted Tyr/Phe parameters were employed (as described above and summarized in Tables S1 and S2). **(a)** Arg–Tyr (parallel; as in the main text). **(b)** Lys–Phe (parallel; as in the main text). **(c)** Arg–Tyr (t-shaped). **(d)** Lys–Phe (t-shaped). Error bars are shown as bands and represent the standard deviations obtained by bootstrapping the results from three independent simulations. **(e)** Variation in the free energy minimum (obtained from the profiles in a–d, with associated error bars) with salt. Upper-case one-letter amino acid codes identify each pair interaction; lower-case p and t denote parallel and t-shaped geometries, respectively.

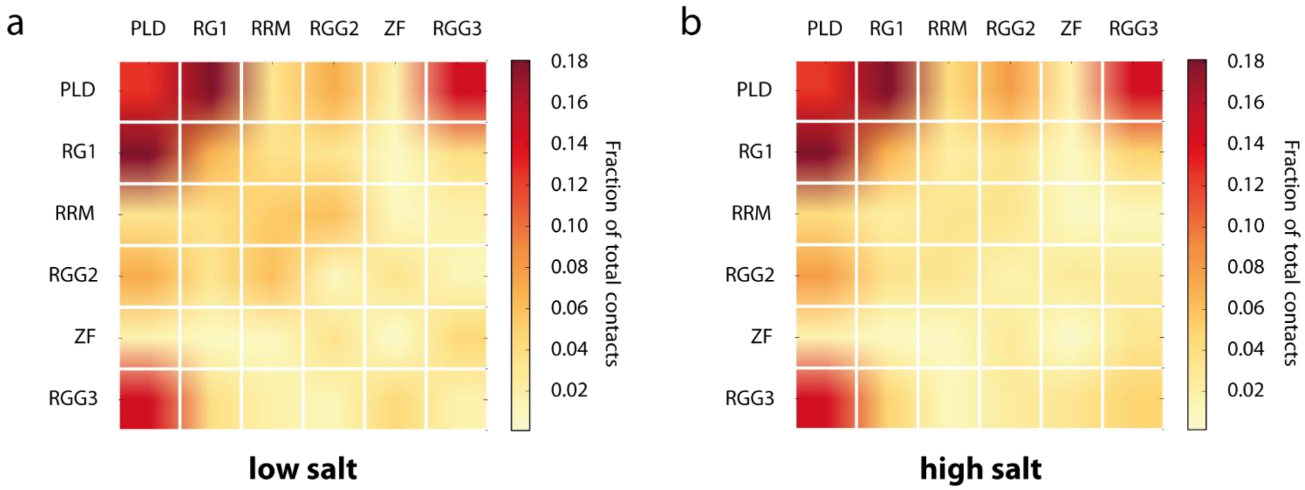


**Figure S4. Frequency of contacts between FUS domains within condensates at (a) low and (b) high salt.** The FUS domains are as follows: prion-like domain (PLD: residues 1–165), arginine–glycine–glycine rich regions (RGG1: residues 166–267; RGG2: residues 371–421; RGG3: residues 454–526), RNA recognition motif (RRM: residues 282–370), and zinc finger (ZF: residues 422–453).

### Supplementary References


1. Patel, A. *et al.* A Liquid-to-Solid Phase Transition of the ALS Protein FUS Accelerated by Disease Mutation. *Cell* **162**, 1066–1077 (2015).
2. Maharana, S. *et al.* RNA buffers the phase separation behavior of prion-like RNA binding proteins. *Science* **360**, 918–921 (2018).
3. Liao, Y. C. *et al.* RNA Granules Hitchhike on Lysosomes for Long-Distance Transport, Using Annexin A11 as a Molecular Tether. *Cell* **179**, 147-164.e20 (2019).
4. Lemaitre, R. P., Bogdanova, A., Borgonovo, B., Woodruff, J. B. & Drechsel, D. N. FlexiBAC: A versatile, open-source baculovirus vector system for protein expression, secretion, and proteolytic processing. *BMC Biotechnol.* **19**, 1–11 (2019).
5. Pronk, S. *et al.* GROMACS 4.5: a high-throughput and highly parallel open source molecular simulation toolkit. *Bioinformatics* **29**, 845–854 (2013).
6. Best, R. B., Zheng, W. & Mittal, J. Balanced Protein–Water Interactions Improve Properties of Disordered Proteins and Non-Specific Protein Association. *J. Chem. Theory Comput.* **10**, 5113–5124 (2014).
7. Benavides, A. L., Aragones, J. L. & Vega, C. Consensus on the solubility of NaCl in water from computer simulations using the chemical potential route. *J. Chem. Phys.* **144**, 124504 (2016).
8. Essmann, U. *et al.* A smooth particle mesh Ewald method. *J. Chem. Phys.* **103**, 8577–8593 (1995).
9. Kumar, S., Rosenberg, J. M., Bouzida, D., Swendsen, R. H. & Kollman, P. A. THE weighted histogram analysis method for free-energy calculations on biomolecules. I. The method. *J. Comput. Chem.* **13**, 1011–1021 (1992).
10. Hub, J. S., De Groot, B. L. & Van Der Spoel, D. G-whams-a free Weighted Histogram Analysis implementation including robust error and autocorrelation estimates. *J. Chem. Theory Comput.* **6**, 3713–3720 (2010).
11. Khan, H. M. *et al.* Improving the Force Field Description of Tyrosine–Choline Cation−π Interactions: QM Investigation of Phenol–N(Me)4+ Interactions. *J. Chem. Theory Comput.* **12**, 5585–5595 (2016).
12. Caldwell, J. W. & Kollman, P. A. Cation-.pi. Interactions: Nonadditive Effects Are Critical in Their Accurate Representation. *J. Am. Chem. Soc.* **117**, 4177–4178 (1995).
13. Frisch, M. J. *et al.* Gaussian 09. Revision D.01, Gaussian, Inc., Wallingford CT (2013).
14. Bayly, C. I., Cieplak, P., Cornell, W. D. & Kollman, P. A. A well-behaved electrostatic potential based method using charge restraints for deriving atomic charges: The RESP model. *J. Phys. Chem.* **97**, 10269–10280 (1993).
15. Dignon, G. L., Zheng, W., Kim, Y. C., Best, R. B. & Mittal, J. Sequence determinants of protein phase behavior from a coarse-grained model. *PLOS Comput. Biol.* **14**, e1005941 (2018).
16. Kapcha, L. H. & Rossky, P. J. A Simple Atomic-Level Hydrophobicity Scale Reveals Protein Interfacial Structure. *J. Mol. Biol.* **426**, 484–498 (2014).
17. Humphrey, W., Dalke, A. & Schulten, K. VMD: Visual molecular dynamics. *J. Mol. Graph.* **14**, 33–38 (1996).
18. Ladd, A. J. C. & Woodcock, L. V. Triple-point coexistence properties of the lennard-jones system. *Chem. Phys. Lett.* **51**, 155–159 (1977).
19. Espinosa, J. R., Sanz, E., Valeriani, C. & Vega, C. On fluid-solid direct coexistence simulations: The pseudo-hard sphere model. *J. Chem. Phys.* **139**, 144502 (2013).
20. García Fernández, R., Abascal, J. L. F. & Vega, C. The melting point of ice Ih for common water models calculated from direct coexistence of the solid-liquid interface. *J. Chem. Phys.* **124**, 144506 (2006).
21. J. S. Rowlinson and B. Widom. *Molecular Theory of Capillarity*. (Clarendon Press, 1984).
22. Plimpton, S. Fast Parallel Algorithms for Short-Range Molecular Dynamics. *J. Comput. Phys.* **117**, 1–19 (1995).
23. Kang, J., Lim, L., Lu, Y. & Song, J. A unified mechanism for LLPS of ALS/FTLD-causing FUS

as well as its modulation by ATP and oligonucleic acids. *PLoS Biol.* **17**, e3000327 (2019).
24. Wang, J. *et al.* A Molecular Grammar Governing the Driving Forces for Phase Separation of Prion-like RNA Binding Proteins. *Cell* **174**, 688–699 (2018).
25. Murthy, A. C. *et al.* Molecular interactions underlying liquid−liquid phase separation of the FUS low-complexity domain. *Nat. Struct. Mol. Biol.* **26**, 637–648 (2019).
26. Qamar, S. *et al.* FUS Phase Separation Is Modulated by a Molecular Chaperone and Methylation of Arginine Cation-π Interactions. *Cell* **173**, 720-734.e15 (2018).
27. Das, S., Lin, Y.-H., Vernon, R. M., Forman-Kay, J. D. & Chan, H. S. Comparative Roles of Charge, Pi, and Hydrophobic Interactions in Sequence-Dependent Phase Separation of Intrinsically Disordered Proteins. *arXiv* 2005.06712 (2020).
28. Gowers, R. *et al.* MDAnalysis: A Python Package for the Rapid Analysis of Molecular Dynamics Simulations. in *Proceedings of the 15th Python in Science Conference* 98–105 (SciPy, 2016). doi:10.25080/majora-629e541a-00e
29. Michaud-Agrawal, N., Denning, E. J., Woolf, T. B. & Beckstein, O. MDAnalysis: A toolkit for the analysis of molecular dynamics simulations. *J. Comput. Chem.* **32**, 2319–2327 (2011).
